# Drug repurposing for mitochondrial diseases using a pharmacological model of complex I deficiency in the yeast *Yarrowia lipolytica*

**DOI:** 10.1101/2020.01.08.899666

**Authors:** Ethan O. Perlstein

**Affiliations:** Perlara PBC, 2625 Alcatraz Ave #435, Berkeley, California 94705, USA

**Keywords:** mitochondrial diseases, Complex I deficiency, drug repurposing, *Yarrowia lipolytica*, Leigh syndrome, estrogens

## Abstract

Mitochondrial diseases affect 1 in 5,000 live births around the world. They are caused by inherited or *de novo* mutations in over 350 nuclear-encoded and mtDNA-encoded genes. There is no approved treatment to stop the progression of any mitochondrial disease despite the enormous global unmet need. Affected families often self-compound cocktails of over-the-counter vitamins and generally recognized as safe nutritional supplements that have not received regulatory approval for efficacy. Finding a new use for an approved drug is called repurposing, an attractive path for mitochondrial diseases because of the reduced safety risks, low costs and fast timelines to a clinic-ready therapy or combination of therapies. Here I describe the first-ever drug repurposing screen for mitochondrial diseases involving complex I deficiency, e.g., Leigh syndrome, using the yeast *Yarrowia lipolytica* as a model system. Unlike the more commonly used yeast *Saccharomyces cerevisiae* but like humans, *Yarrowia lipolytica* has a functional and metabolically integrated respiratory complex I and is an obligate aerobe. In 384-well-plate liquid culture format without shaking, *Yarrowia lipolytica* cells grown in either glucose-containing media or acetate-containing media were treated with a half-maximal inhibitory concentration (3µM and 6μM, respectively) of the natural product and complex I inhibitor piericidin A. Out of 2,560 compounds in the Microsource Spectrum collection, 24 suppressors of piercidin A reached statistical significance in one or both media conditions. The suppressors include calcium channel blockers nisoldipine, amiodarone and tetrandrine as well as the farnesol-like sesquiterpenoids parthenolide, nerolidol and bisabolol, which may all be modulating mitochondrial calcium homeostasis. Estradiols and synthetic estrogen receptor agonists are the largest class of suppressors that rescue growth of piericidin-A-treated *Yarrowia lipolytica* cells in both glucose-containing and acetate-containing media. Analysis of structure-activity relationships suggests that estrogens may enhance bioenergetics by evolutionarily conserved interactions with mitochondrial membranes that promote mitochondrial filamentation and mitochondrial DNA replication.

## Introduction

Mitochondrial diseases are genetically heterogenous, multi-system metabolic disorders affecting both children and adults in all populations across the globe (Maldonado *et al.*, 2019). There are many types of mitochondrial diseases depending on which gene is mutated and which component of the oxidative phosphorylation (OXPHOS) pathway is proximally affected. Most mitochondrial disease genes are encoded in the nucleus, but some are among the 13 protein-coding genes and 24 tRNA and rRNA genes encoded by the 16,569 base pair mitochondrial genome, or mtDNA. Mitochondrial disease mutations can be either inherited or *de novo*. The fraction of mutated mtDNA in a cell or person is referred to as heteroplasmy, which can vary from tissue to tissue and over the life of a patient starting in infancy. The evolutionary conservation of mitochondrial disease genes combined with the phenotypic variability of clinical presentations present challenges to creating therapeutically relevant disease models that are also amenable to high-throughput drug screens, genetic modifier screens, and biomarker discovery (Lasserre *et al.*, 2015; Maglioni & Ventura, 2016; Dancy *et al.*, 2015; Sen & Cox, 2017).

Yeast, and more broadly fungi, are single-celled animals whose mitochondrial biology is exquisitely conserved. They serve as the logical starting point for a cross-species disease modeling and drug screening approach (Malina *et al.*, 2018; Awad *et al.*, 2018; Lao *et al.*, 2019). The most commonly used yeast species in the lab and a foundational model organism, *Saccharomyces cerevisiae* (budding yeast), has been repeatedly validated as a relevant model for mitochondrial diseases (Sesaki *et al.*, 2014; Kaliszewska *et al.*, 2015; Heimer *et al.*, 2016). The most modeled mitochondrial disease gene in yeast is *SURF1*: a nuclear-encoded assembly factor subunit for complex V in which recessive loss-of-function mutations cause Leigh syndrome, and whose ortholog in *Saccharomyces cerevisiae* is called *SHY1* (Zeviani *et al.*, 1999; Barrientos *et al.*, 2002; Reinhold *et al.*, 2011). In those studies, it was shown that human *SURF1* can replace the function of yeast *SHY1*, but high-throughput drug screens were not attempted. Two advantages of yeast disease models are affordability and rapidity of drug screening based on a simple “growth/no growth” phenotypic readout. The first drug repurposing screen for mitochondrial disease in *Saccharomyces cerevisiae* was published in 2014 using a knockout of a complex V assembly factor gene, but the two most promising candidates from the screen did not have a clear path to clinical development and so further work was discontinued (Aiyar *et al.*, 2014).

There are two significant disadvantages to using *Saccharomyces cerevisiae* as a human mitochondrial disease model. First, *Saccharomyces cerevisiae* lacks a respiratory complex I, unlike other fungi (Matus-Ortega *et al.*, 2015). Therefore, the coordinated expression and activity of complex I subunits with each other and with downstream electron transport chain (ETC) complexes and metabolic pathways cannot be modeled in *Saccharomyces cerevisiae*. Second, *Saccharomyces cerevisiae* is a facultative aerobe which can survive completely lacking mitochondria as so called “petites” – unlike in higher eukaryotes. In an attempt to address those deficits, another yeast species called *Schizosaccharomyces pombe* (fission yeast), which does express a respiratory complex I, has been employed as a model system for mitochondrial diseases. In early 2019, a group in France published a 1,760-compound drug repurposing screen using a *Schizosaccharomyces pombe* genetic model of mitochondrial disease caused by mutations in *OPA1*, a mitochondrial dynamin-like protein that regulates mitochondrial fusion and whose yeast ortholog is called *MSP1* (Delerue *et al.*, 2019). They discovered that hexestrol, an estrogen mimic and synthetic estrogen receptor agonist, has a mitoprotective mechanism of action. The mitochondrial fragmentation phenotype and loss of mtDNA nucleotids phenotype exhibited by *msp1*^*P300S*^ mutant *Schizosaccharomyces pombe* cells were both completely suppressed by hexestrol. In fact, Delerue *et al* showed that hexestrol stimulated mitochondrial hyperfilamentation and increase mtDNA nucleotid copy number even in wildtype cells.

Encouraged by those results, I decided to conduct a similar drug repurposing screen but with a larger drug repurposing library, two different growth media conditions, a pharmacological model instead of a genetic model, and the yeast *Yarrowia lipolytica* instead of *Schizosaccharomyces pombe*. As of this submission, a PubMed search using terms “*Yarrowia lipolytica*” and “mitochondria” resulted in 103 publications, while the analogous search for “*Saccharomyces cerevisiae*” and “mitochondria” yielded over 8,400 publications (and “*Schizosaccharomyces pombe*” and “mitochondria” revealed 353 publications). In spite of the comparatively minuscule number of publications and small number of mitochondria labs using it as a model system, *Yarrowia lipolytica* has been recognized, and is gaining wider acceptance, as a model system for mitochondrial biology. For example, the crystal structure of electron transport chain complexes was recently solved using *Yarrowia lipolytica* proteins (Zickermann *et al.*, 2015; Hahn *et al.*, 2016), and more recently a cryo-EM structure of complex I from *Yarrowia lipolytica* in action was published (Parey *et al.*, 2018).

I decided to follow the drug repurposing template created at Perlara PBC, the first biotech public benefit corporation focused on rare genetic diseases, for the congenital disorder of glycosylation PMM2-CDG (Iyer *et al.*, 2019). That effort resulted in the discovery of the generic Japanese diabetic neuropathy drug epalrestat as the first-in-class PMM2 enzyme activator, which is currently in a single patient clinical study. Over the course of three months of experiment time and for a cost of goods, labor and overhead of approximately $33,000, I performed the first-ever drug repurposing screen of *Yarrowia lipolytica* cells treated with the natural product and complex I inhibitor piericidin A (Zhou & Fenical, 2016). 24 piercidin A suppressors were identified spanning different mechanisms of action and pharmacological classes, including the unbiased and independent discovery of the synthetic estrogen receptor agonist hexestrol, which was the top hit from the *Schizosaccharomyces pombe* drug repurposing screen (Delerue *et al.*, 2019). Structure-activity relationship (SAR) analysis and a counter-screen for rapamycin suppressors demonstrated the specificity of piercidin A suppressors. Piericidin A suppressors are repurposing candidates poised for validation studies in mitochondrial disease patient-derived cells (fibroblasts or iPSCs) and in more biologically complex animals models of mitochondrial disease (Soma *et al.*, 2018).

## Materials and Methods

### Strains, growth conditions and compounds

The *Yarrowia lipolytica* wildtype strain PO1f was used in drug repurposing screens and was purchased from ATCC (MYA-2613). Screening was conducted using the 2,560-compound Microsource Spectrum collection consisting of FDA approved drugs, bioactive tool compounds, and plant-based natural products (Microsource Discovery Systems, Inc). W303-1a was used in the rapamycin counter-screen and a MAT**a** prototrophic version of the *Saccharomyces cerevisiae* S288C strain was used in the rapamycin suppressor dose-response retests; both strains were generously provided by Dr. Maitreya Dunham from the University of Washington. YPD (2% glucose, 1% yeast extract and 2% peptone) or YPA (2% acetate, 1% yeast extract and 2% peptone) was buffered to pH 6.5 with HEPES and autoclaved before use. Greiner Bio-One polystyrene clear-bottom 384-well plates were purchased from Sigma-Aldrich. A SpectraMax plate reader (Molecular Devices, Inc) was used for all OD_600_ absorbance measurements. Piercidin A was purchased from Cayman Chemical. Rapamycin was purchased from Sigma-Aldrich. Rapamycin suppressors were also purchased from Sigma-Aldrich. Dose-response experiments were performed as previously described (Lao *et al.*, 2019).

### High-throughput 384-well-plate growth assay in yeast

125nL of test compound from the Microsource Spectrum library (or DMSO in the case of control wells) was acoustically dispensed into each well of a 384-well plate using an Echo550 acoustic dispenser (manufactured by Labcyte, Inc., which was acquired by Beckman Coulter in 2019). These plates were stored at −80°C until use. Overnight cultures of *Yarrowia lipolytica* or *Saccharomyces cerevisiae* were grown in 2mL of YPD (or YPA where appropriate) and then diluted 1:1000 into fresh media. 50μL of media with yeast and primary compound (piericidin A or rapamycin) was manually dispensed using a 16-channel pipettor into the 384-well plates which had been pre-dispensed with test compounds. Plates were covered using Ambryx optically clear breathable plate seals and then incubated on the benchtop for up to five days. Plates were vortexed for 30 seconds at maximum speed in order to resuspend the yeast growing at the bottom of each well, and then analyzed by the plate reader.

### Z-score analysis and structure-activity relationships

Raw OD_600_ absorbance measurements were exported as .csv files from the SpectraMax software package. Absorbance measurements for each condition were converted to Z-scores and then rank ordered for analysis. Chemical structures were rendered in ChemDraw. Raw and processed datasets are available in Supplementary Materials.

## Results

### Drug repurposing screen using a pharmacological model of complex I deficiency

Based on the results of dose-response experiments, the complex I inhibitor piericidin A (**Figure 1A**) and ~0.5 × 10^6^ *Yarrowia lipolytica* cells suspended in either glucose-containing media (hereafter YPD) with 3μM piericidin A or acetate-containing media (hereafter YPA) with 6μM piericidin A were dispensed in duplicate into clear-bottom 384-well plates. Columns 1, 2, 23, 24 are DMSO controls, i.e., piericidin-A-treated cells with DMSO vehicle. Columns 3 to 22 and rows A to P contained a droplet of acoustically dispensed test compound from the Microsource Spectrum collection such that addition of 50μL of fresh media containing yeast cells and an IC_50_ dose of piericidin A yielded a final test compound concentration of 25μM. Plates were incubated without shaking for up to five days with optical density measurements taken daily in a plate reader. YPD plates were subjected to absorbance measurements at OD_600_ in a plate reader.

**Figure 1.**
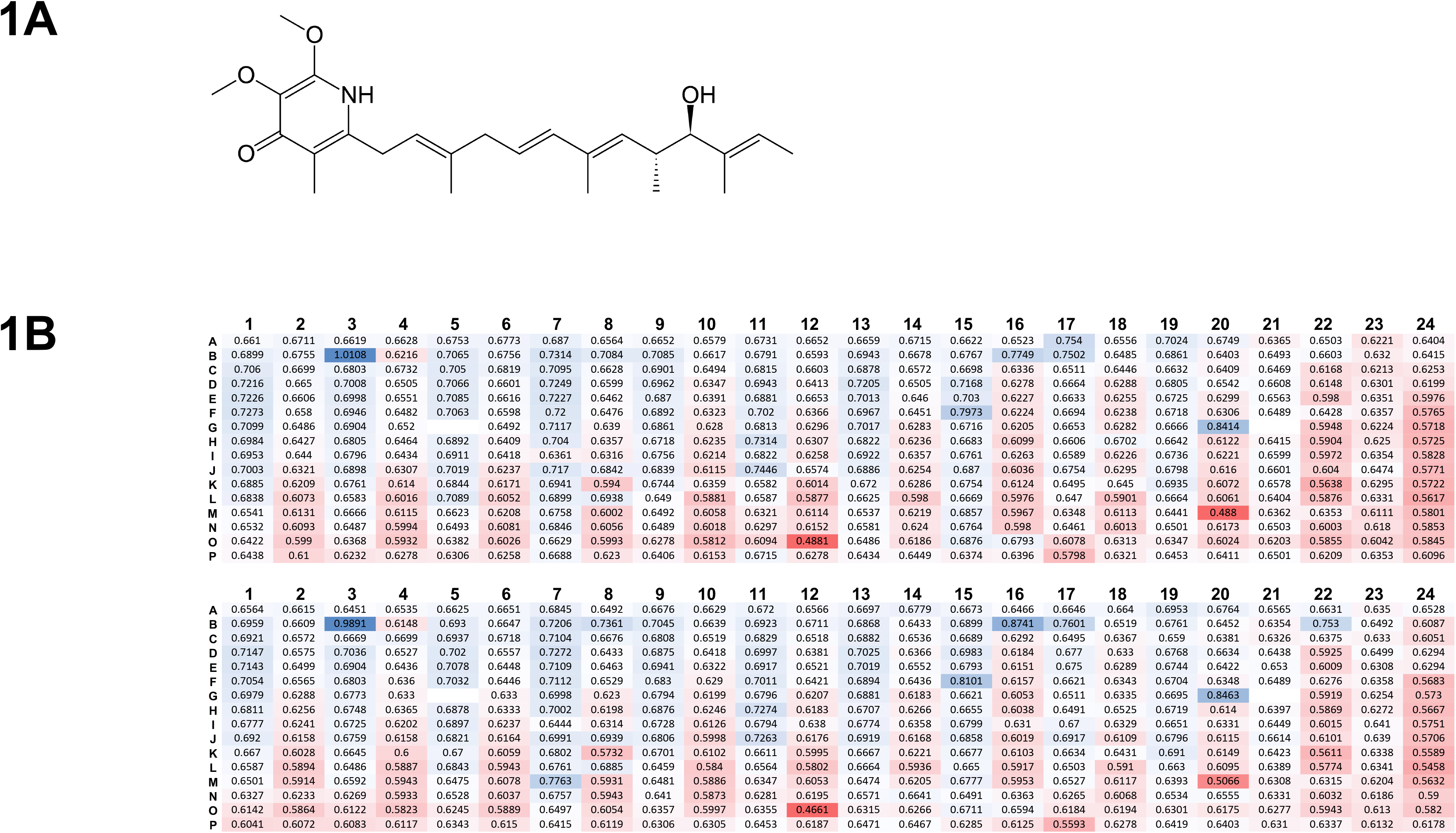
Overview of complex I deficiency drug repurposing screen. (A) The chemical structure of piericidin A. (B) Two replicates of a representative plate from the screen. Columns and rows are labeled. Values are raw OD_600_ absorbance measurements. Columns 1, 2, 23 and 24 are DMSO controls. High OD_600_ values are in blue while low OD_600_ values are red. An example of a presumptive suppressor is B03. An example of a presumptive enhancer is O12. Wells G05 and G21 show no value because these wells contained darkly hued compounds with artificially high OD_600_ values.

There were 96 conditions in total: three timepoints × eight library plates × two replicates × two growth conditions. A representative plate is shown in **Figure 1B**. OD_600_ measurements were converted to Z-scores for rank-order analysis and structure-activity relationships (SAR) analysis. Replicate one and replicate two were plotted against each other to determine statistically significant hits in both replicates for further consideration. There was better agreement between YPA plate replicate pairs versus YPD plate replicate pairs. Z-score plots are shown in **Figure 2**. 24 statistically significant suppressors of piericidin A are listed in **Table 1** and summarized below.

**Table 1.** 24 piericidin A suppressors.

| Condition | Suppressors with Z > 5 in both replicates | Suppressors with Z > 4 in at least one replicate |
| --- | --- | --- |
| YPD 48 hours | 1<br>**totarol-19-carboxylic acid, methyl ester** | 5<br>**<u>dienestrol</u>**; **totarol**; melengestrol; celastrol; **7-desacetoxy-6,7-dehydrogedunin** |
| YPD 64 hours | 0 | 0 |
| YPD 84 hours | 2<br><u>prochlorperazine</u>; **<u>promethazine</u>** | 2<br><u>exemestane</u>; oxolamine |
| YPA 64 hours | 4<br><u>nisoldipine</u>; **7-desacetoxy-6,7-dehydrogedunin**; **totarol-19-carboxylic acid, methyl ester**; **totarol** | 4<br>bisabolol; cedrelone; nerolidol; eugenyl benzoate |
| YPA 84 hours | 8<br><u>nisoldipine</u>; **7-desacetoxy-6,7-dehydrogedunin**; **totarol-19-carboxylic acid, methyl ester**; **totarol**; bisabolol; nerolidol; eugenyl benzoate; cedrelone | 8<br>tetrandrine; **<u>dienestrol</u>**; robustic acid; dihydrocelastrol; dihydrogedunin; <u>amiodarone</u>; parthenolide; <u>naftifine</u> |
| YPA 108 hours | 9<br><u>nisoldipine</u>; **7-desacetoxy-6,7-dehydrogedunin**; **totarol-19-carboxylic acid, methyl ester**; cedrelone; dihydrogedunin; eugenyl benzoate; nerolidol; **<u>promethazine</u>**; tomatidine | 6<br>**totarol**; bisabolol; dihydrocelastrol; robustic acid; acetylsalicylanilide; parthenolide |

*Compounds in bold are hits in both YPD and YPA. Underlined compounds are FDA approved.*

**Table 2.** Project cost summary.

| Activity | Cost (\$) |
| --- | --- |
| 1 FTE for 3 months | 15000 |
| Bench space for 3 months | 3000 |
| Echo550 liquid dispensing for 200 plates | 6580 |
| Microsource Spectrum library (1µL) | 1500 |
| 200 384-well plates | 1000 |
| 400 plates seals | 750 |
| Compounds | 3000 |
| Other reagents and consumables | 1500 |
| Total | **32330** |

**Figure 2.**
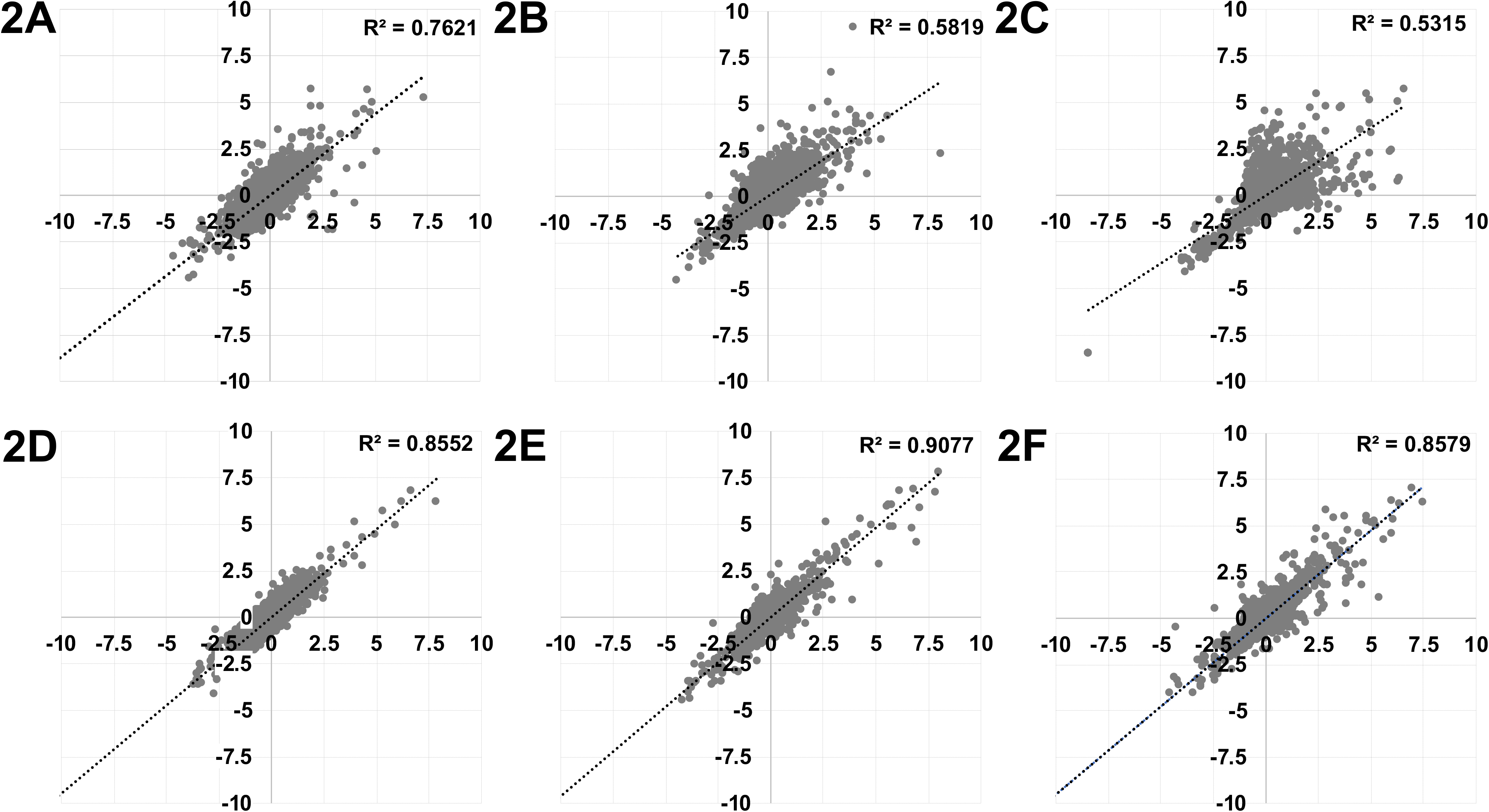
Z-score summary plots of complex I deficiency drug repurposing screen. Z-scores of each test compound (gray circles) from replicate one and replicate two were plotted against each other. X-axis and y-axis values are Z-scores. Dotted black line indicates linear regression with r-squared values shown in the upper right quadrant. (A) YPD at the 48-hour timepoint. (B) YPD at the 64-hour timepoint. (C) YPD at the 84-hour timepoint. (D) YPA at the 64-hour timepoint. (E) YPA at the 84-hour timepoint. (F) YPA at the 108-hour timepoint.

### Novel piericidin A suppressors comprise multiple mechanistic classes

Using a stringent threshold for significance, YPD plates at the 48-hour timepoint yielded only one test compound with a Z-score greater than five in both replicates: totarol-19-carboxylic acid, methyl ester. When the threshold for significance is relaxed to test compounds with a Z-score of at least four in replicate one and at least 2.5 in replicate two, there are five suppressors: dienestrol; totarol; melengestrol; celastrol; 7-desacetoxy-6,7-deydrogedunin. There are no test compounds that pass the stringent or relaxed threshold YPD plates at the 64-hour timepoint. At the 84-hour and final timepoint, YPD plates yielded two test compounds at the stringent threshold: prochlorperazine and promethazine. Two test compounds passed the relaxed threshold at 84 hours in YPD: exemestane and oxolamine. In total, 10 test compounds pass one or both piericidin A suppressor thresholds in the YPD condition.

5/10 (50%) of the above piericidin A suppressors in YPD are also piericidin A suppressors in YPA: totarol-19-carboxylic acid, methyl ester; totarol; dienestrol; 7-acetoxy-6,7-dehydrogedunin. Overall, the hit rate is roughly double in the YPA condition than in the YPD condition (0.74% versus 0.39%). Using a stringent threshold for significance, YPA plates at the 84-hour timepoint yielded four test compounds with a Z-score greater than five in both replicates: nisoldipine; 7-acetoxy-6,7-dehydrogedunin; totarol-19-carboxylic acid, methyl ester; totarol. When the threshold for significance is relaxed to test compounds with a Z-score of at least four in replicate one and at least 2.5 in replicate two, there are four piericidin A suppressors: bisabolol, cedrelone, nerolidol and eugenyl benzoate.

YPA plates at the 84-hour timepoint yielded eight test compounds that pass the stringent threshold: nisoldipine; 7-acetoxy-6,7-dehydrogedunin; totarol-19-carboxylic acid, methyl ester; totarol; bisabolol; nerolidol; eugenyl benzoate; cedrelone. At 84 hours in YPA, eight test compounds pass the relaxed threshold: tetrandrine; dienestrol; robustic acid; dihydryocelastrol; dihydrogedunin; amiodarone; parthenolide; naftifine. At the 108-hour and final timepoint, YPA plates yielded nine test compounds at the stringent threshold: nisoldipine; 7-acetoxy-6,7-dehydrogedunin; totarol-19-carboxylic acid, methyl ester; cedrelone; dihydrogedunin; eugenyl benzoate; nerolidol; promethazine; tomatidine. Six test compounds pass the relaxed threshold: totarol; bisabolol; dihydrocelastrol; robustic acid; acetylsalicylanilide; parthenolide. In total, 19 test compounds pass one or both piericidin A suppressor thresholds in YPA.

Six compounds are highlighted in **Figure 3** and **Figure 4** to illustrate three features of piericidin A suppressors: 1) potency, i.e., strong versus modest suppression; 2) onset, i.e., fast-acting versus slow-acting suppression; 3) conditionality, i.e., suppression in one or both media conditions. At the early timepoint (48 hours for YPD plates and 64 hours for YPA plates), the strongest and fastest acting piericidin A suppressor is totarol-19-carboxylic acid, methyl ester (**Figures 3A**, **4A**). Nisoldipine is the strongest or second strongest and fastest acting piericidin A suppressor in YPA at all three timepoints. In YPD, nisoldipine is also fast-acting but confers modest suppression. Estradiol and dienestrol are fast-acting piericidin A suppressors but not as strong as the totarols or nisoldipine. Prochlorperazine and promethazine are the strongest piericidin A suppressors in YPD but also the slowest acting. In YPA, prochlorperazine and promethazine appear to have an even slower onset of suppression and more modest potency (**Figures 3C**, **4C**).

**Figure 3.**
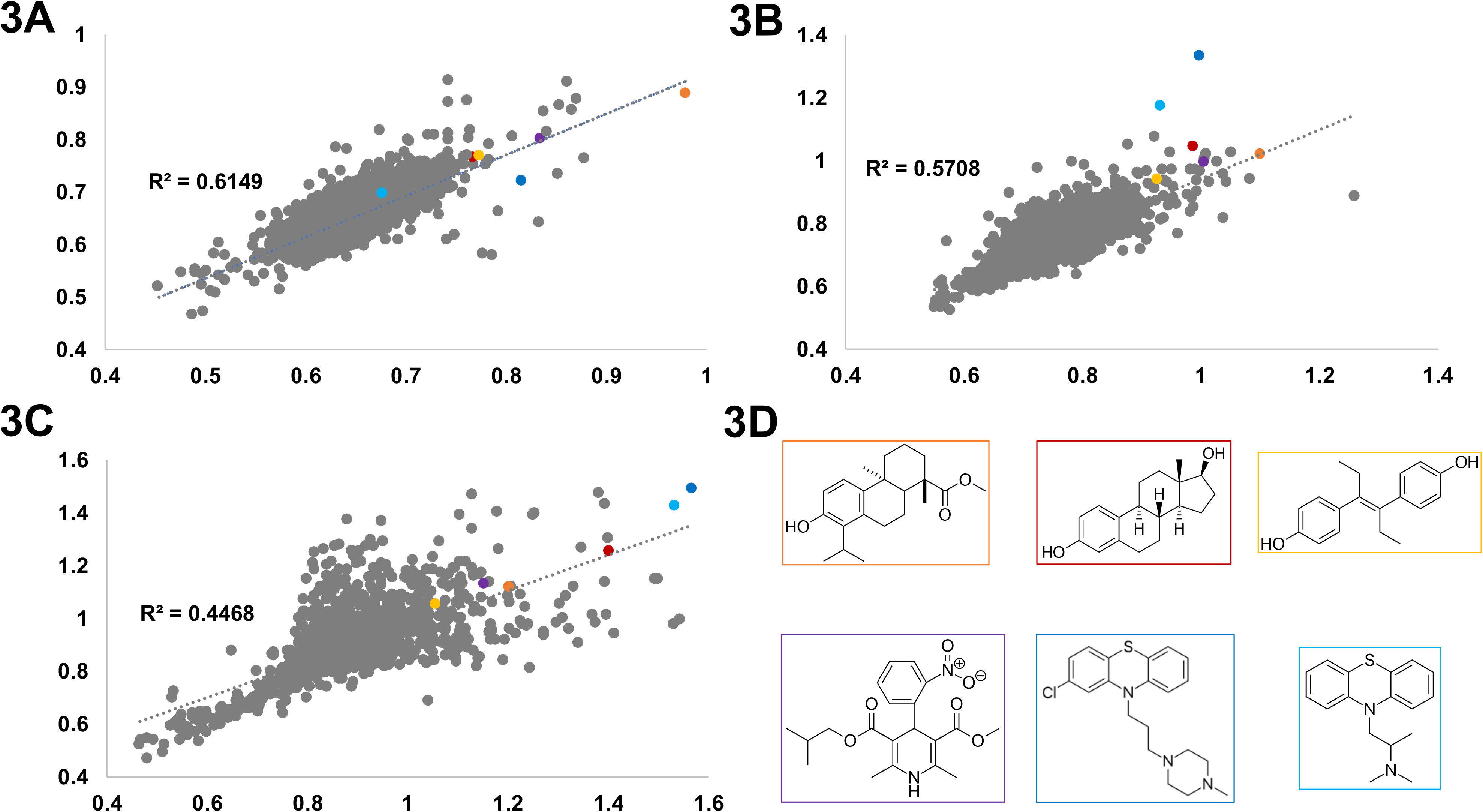
Top piericidin A suppressors in YPD. Replicate OD_600_ absorbance measurements (gray circles) of each test compound in the YPD growth condition were plotted against each other. Dotted black line indicates linear regression with r-squared values shown. Colored circles indicate highlighted suppressors. (A) YPD at the 48-hour timepoint. (B) YPD at the 64-hour timepoint. (C) YPD at the 84-hour timepoint. (D) Chemical structures of six suppressors: totarol-19-carboxylic acid, methyl ester (orange box); estradiol (red box); dienestrol (yellow box); nisoldipine (purple box); prochlorperazine (blue box); promethazine (cyan box).

**Figure 4.**
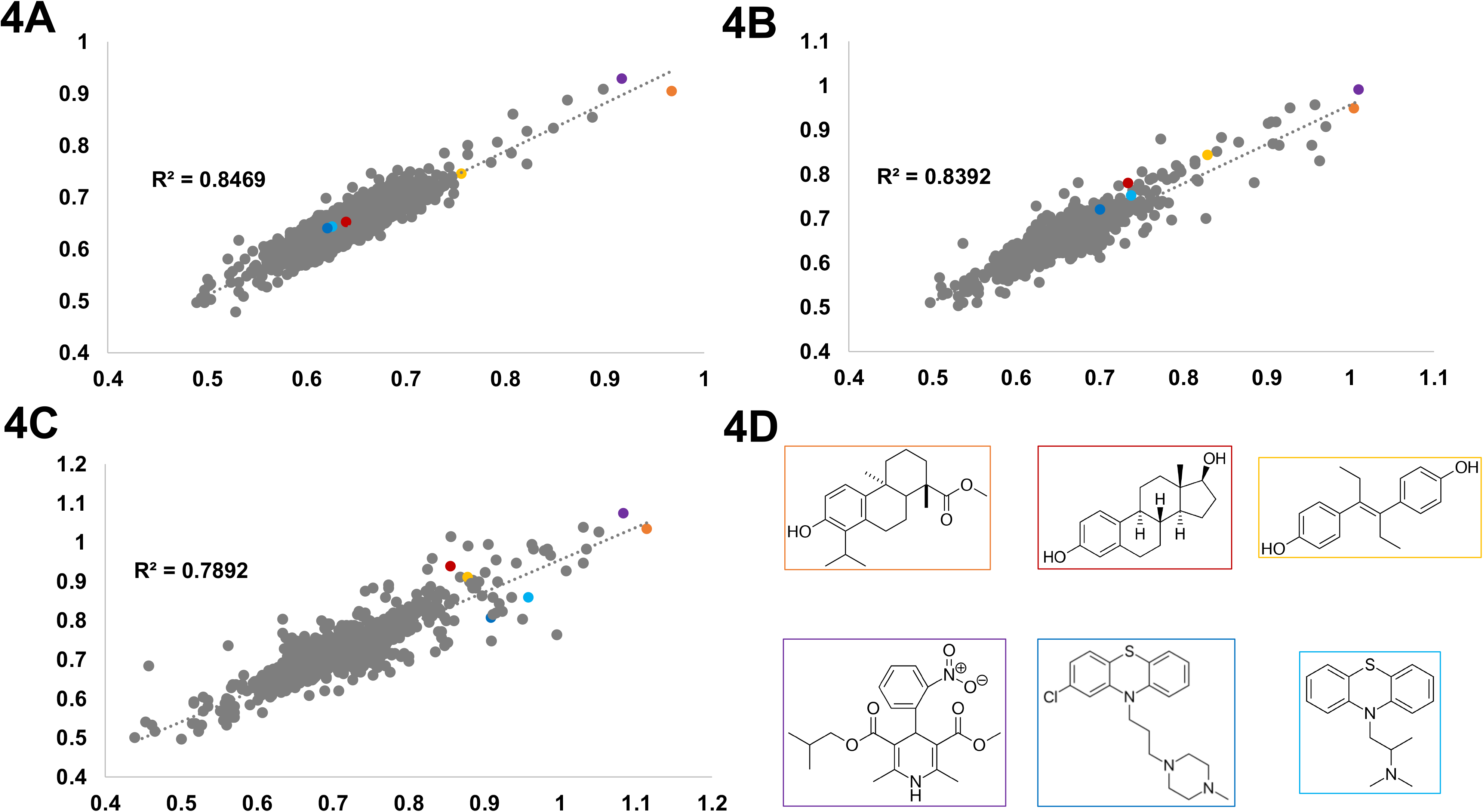
Top piericidin A suppressors in YPA. Replicate OD_600_ absorbance measurements (gray circles) of each test compound in the YPA growth condition were plotted against each other. Dotted black line indicates linear regression with r-squared values shown. Colored circles indicate highlighted suppressors. (A) YPA at the 64-hour timepoint. (B) YPD at the 84-hour timepoint. (C) YPD at the 108-hour timepoint. (D) Chemical structures of six suppressors: totarol-19-carboxylic acid, methyl ester (orange box); estradiol (red box); dienestrol (yellow box); nisoldipine (purple box); prochlorperazine (blue box); promethazine (cyan box).

### Structure-activity relationships of repurposing candidates

The Microsource Spectrum collection is well suited for SAR analysis because it contains most if not all members of many pharmacological classes, including the piericidin A suppressors listed in **Table 1**. The high quality of the primary screening dataset invited SAR analysis even though the screen was performed only at a single concentration of test compound (25μM). Totarol-19-carboxylic acid, methyl ester and the parent compound totarol are phenolic diterpenes and plant-based natural products (Evans *et al.*, 1999). Both totarols are structurally related to the cholesterol-derived steroid estrogen, which is consistent with the fact that half a dozen estrogens are piercidin A suppressors as well. It is also consistent with the fact that phenolic diterpenes have been shown to bind to estrogen receptors (Chun *et al.*, 2014). In particular, estradiol and dienestrol are suppressors across all timepoints and in both media conditions. These results suggested a closer examination of structurally related estrogens (**Figure 5**) and other estrogen receptor agonists, as well as estrogen receptor antagonists (**Figure 6**), in order to ascertain the active pharmacophore.

**Figure 5.**
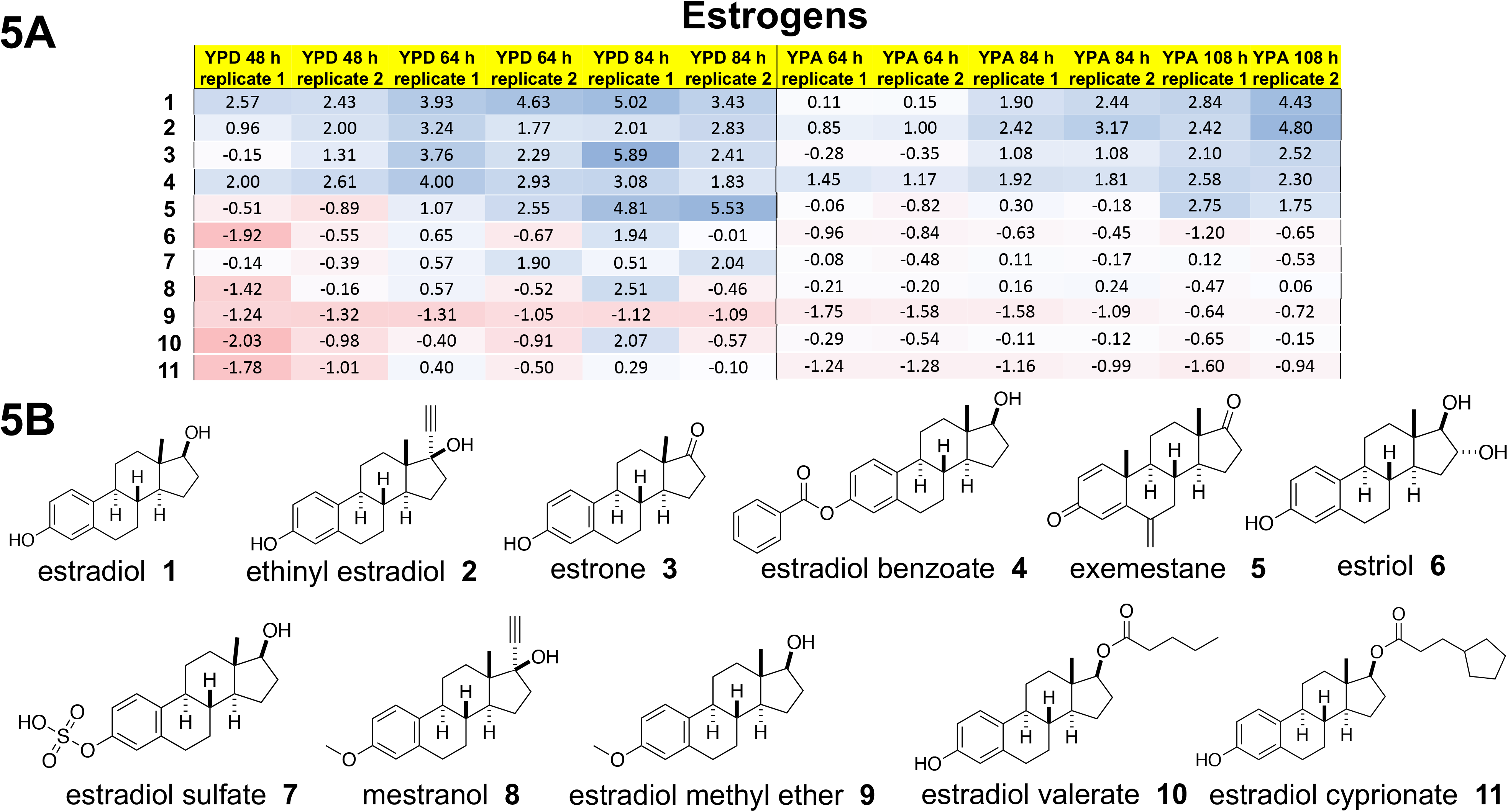
Estrogens are piericidin A suppressors. (A) Heat map of Z-scores. The higher the Z-score, the bluer the cell. The lower the Z-score, the redder the cell. A Z-score of zero is a white cell. Columns are screening conditions and timepoints of absorbance measurements. Rows are compounds. (B) Chemical structures showing structure-activity relationships.

**Figure 6.**
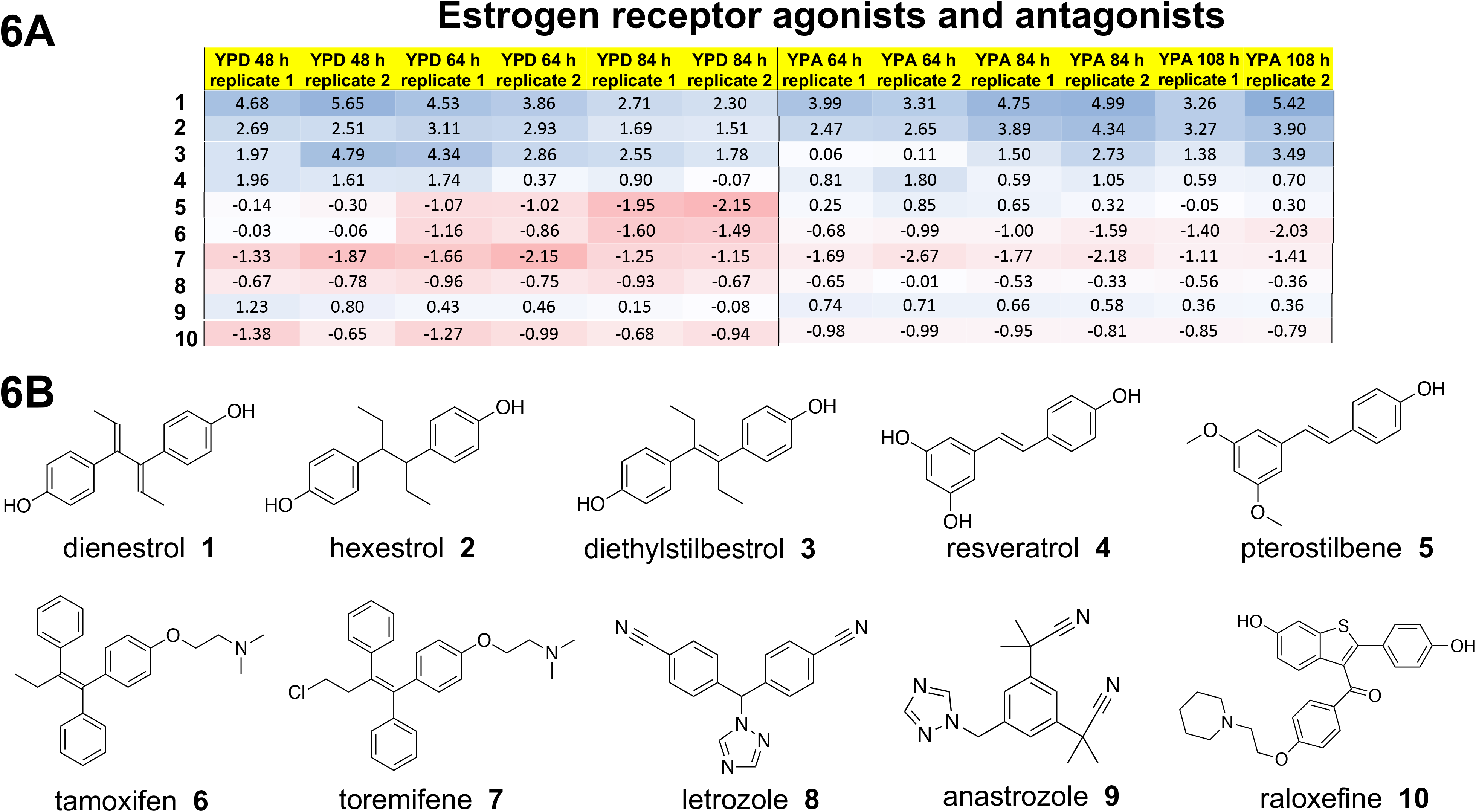
Estrogen receptor modulators are piericidin A suppressors. (A) Heat map of Z-scores. The higher the Z-score, the bluer the cell. The lower the Z-score, the redder the cell. A Z-score of zero is a white cell. Columns are screening conditions and timepoints of absorbance measurements. Rows are compounds. (B) Chemical structures showing structure-activity relationships.

Estradiol and single-substitution analogs like ethinyl estradiol, estrone and estradiol benzoate are approximately equally potent piericidin A suppressors (**Figure 5A**). However, other single-substitution analogs of estradiol like estriol, estradiol sulfate, mestranol, estradiol methyl ether, estradiol valerate and estradiol cyprionate are inactive at all timepoints and in both media conditions (**Figure 5A**). These results suggest the contours of an estrogenic pharmacophore. Substitution of the phenolic hydroxyl is tolerated in the case of estradiol benzoate but not estradiol methyl ether or estradiol sulfate (**Figure 5B**). However, the 17-beta hydroxy group does not tolerate substitutions as shown by the inactivity of estradiol valerate and estradiol cyprionate (**Figure 5B**).

Dienestrol is synthetic estrogen receptor agonist whose structure is based on an open conformation of estradiol (Duax *et al.*, 1985). Hexestrol is a close analog of dienestrol and is an equipotent piericidin A suppressor; diethylstilbestrol is another close analog of dienestrol but it appears less potent (**Figure 6A**). However, resveratrol and pterostilbene, which lack diethyl substitutions, are inactive. The following estrogen receptor antagonists belonging to three different structural classes are all inactive: tamoxifen, toremifene, letrozole, anastrozole and raloxefine (**Figure 6A**). In fact, it appears that toremifene may enhance the cytotoxic effects of piericidin A. Other steroids present in the Microsource Spectrum library (**Figure S1**) that are inactive at 25μM include: cholesterol, androsterone and progesterone. Curiously, the steroid intermediate lanosterol has intermediate potency but only in YPA and the later timepoint.

Nisoldipine is a 1,4-dihydropyridine calcium channel blocker first approved for the treatment of hypertension in 1995 (Plosker & Faulds, 1996). Nine other dihydropyridine calcium channel blockers are present in the Microsource Spectrum collection and their SAR is presented in **Figure 7**. As mentioned above, nisoldipine is one of the top two strongest and fastest acting piericidin A suppressors (along with totarol-19-carboxylic acid, methyl ester) to emerge from this drug repurposing screen. Manidipine appears to be equipotent to nisoldipine in YPD, but in YPA it has modest potency (**Figure 7A**). Nivaldipine and nifedipine are weaker piericidin A suppressors than manidipine. Nifedipine is the closest structural analog to nisoldipine out of the nine dihydropyridines screened, with an additional isopropyl substitution. Conversely, the following five dihydropyridine calcium channel blockers are inactive: nitrendipine; nimodipine; nicardipine; cilnidipine; felodipine; amlodipine. Interestingly, the non-dihydropyridine calcium channel blockers amiodarone, its analog benzbromarone and tetrandrine are also piericidin A suppressors, and like nisoldipine they appear to be active only in YPA (**Figure S2**). In fact, amiodarone appears to be an enhancer of piericidin A cytotoxicity. However, the non-dihydropyridine calcium channel blockers verapamil and diltiazem are inactive.

**Figure 7.**
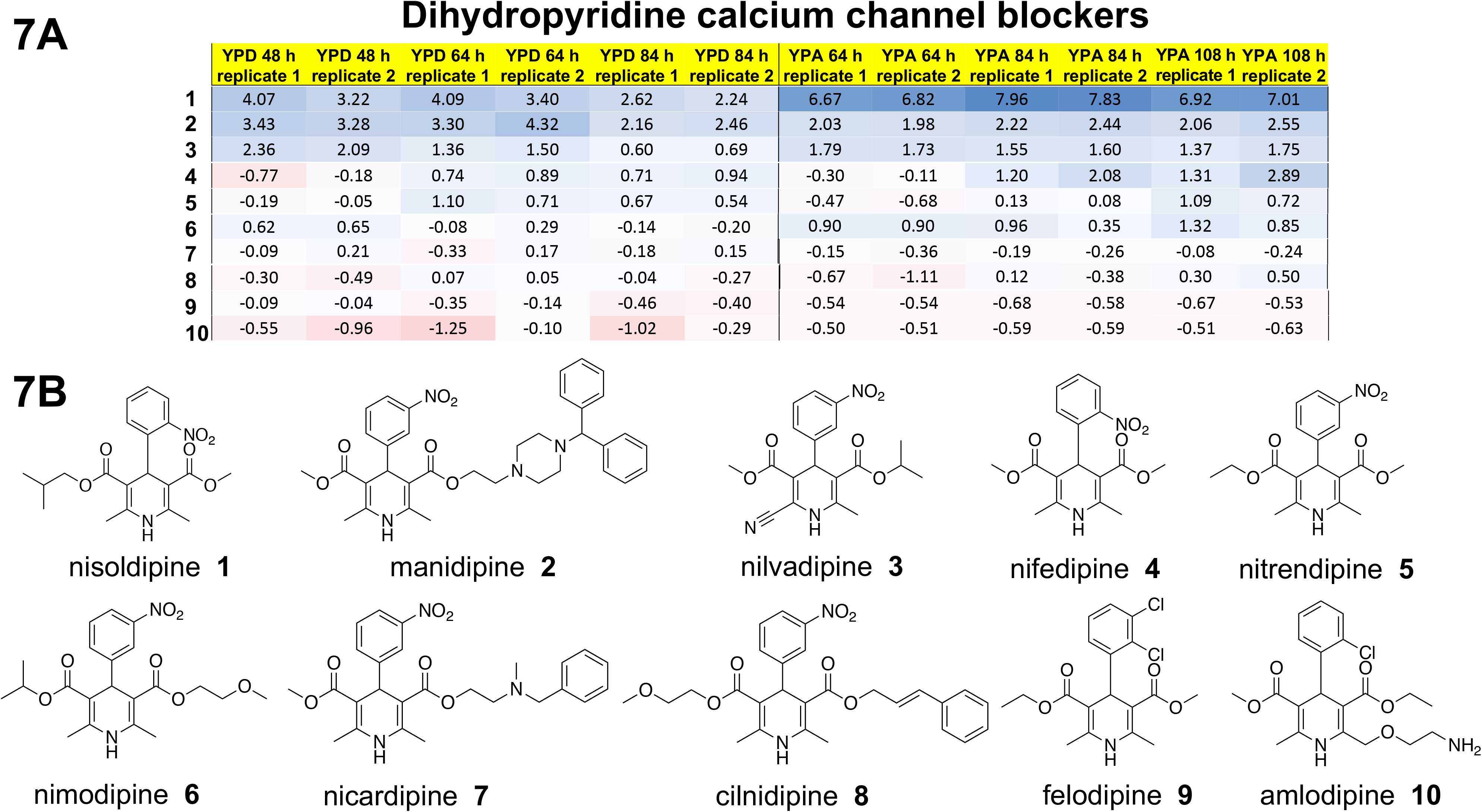
1,4-dihydropyridine calcium channel blockers are piericidin A suppressors. (A) Heat map of Z-scores. The higher the Z-score, the bluer the cell. The lower the Z-score, the redder the cell. A Z-score of zero is a white cell. Columns are screening conditions and timepoints of absorbance measurements. Rows are compounds. (B) Chemical structures showing structure-activity relationships.

As shown in **Figure 8**, the psychoactive drugs prochlorperazine and promethazine are the most potent piericidin A suppressors in the phenothiazine class. Ethopropazine is the next most potent piericidin A suppressor, which is not surprising given that the entire structure of promethazine is contained within ethopropazine. Chlorpromazine has marginal potency. The following phenothiazines are inactive: promazine; trifluoperazine; trifluopromazine; acepromazine. The building blocks phenothiazine and piperazine are also inactive. Phenothiazines are notorious for polypharmacology so it is unclear which target engagement is responsible for piericidin A suppression (Caldara *et al*, 2017).

**Figure 8.**
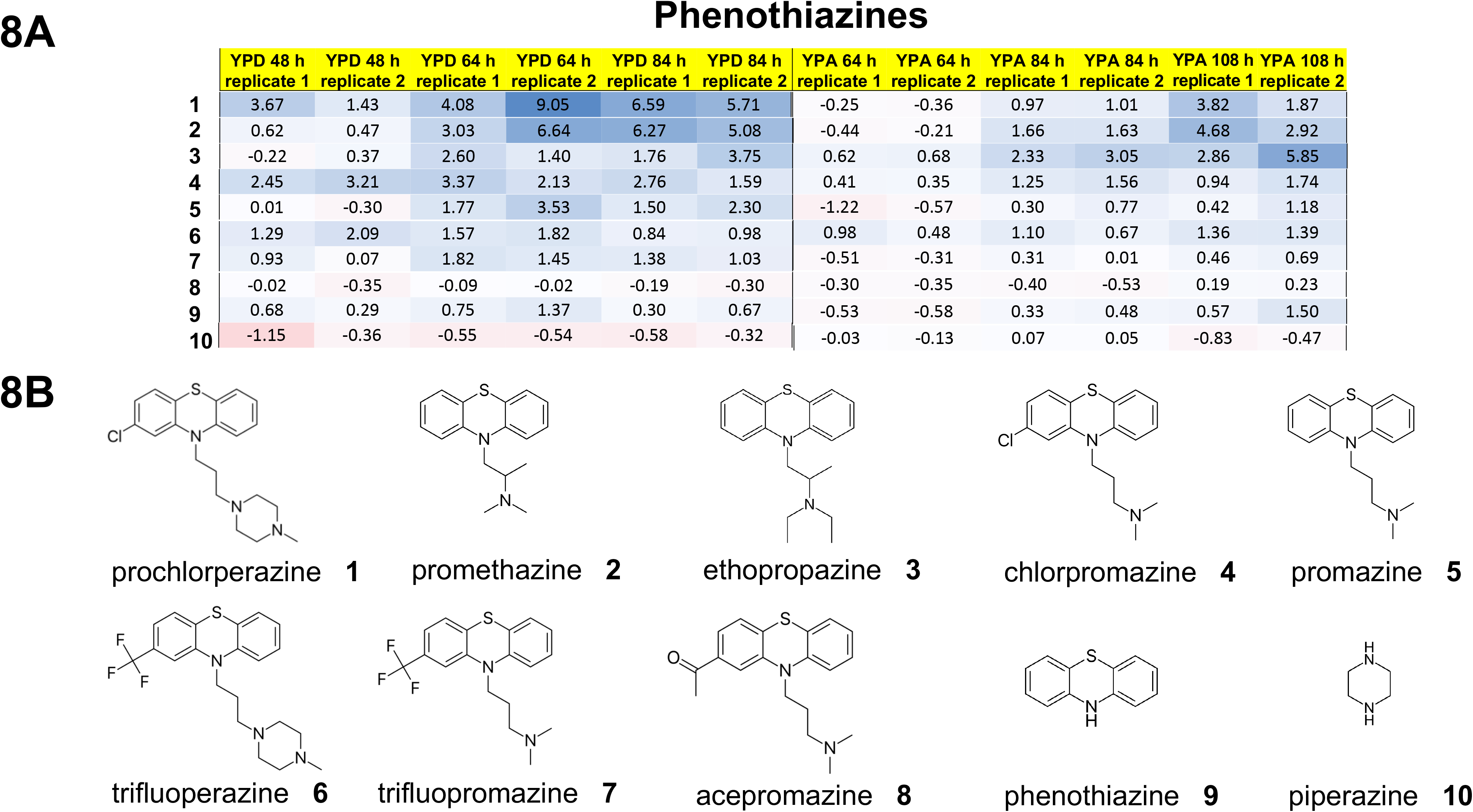
Phenothiazines are piericidin A suppressors. (A) Heat map of Z-scores. The higher the Z-score, the bluer the cell. The lower the Z-score, the redder the cell. A Z-score of zero is a white cell. Columns are screening conditions and timepoints of absorbance measurements. Rows are compounds. (B) Chemical structures showing structure-activity relationships.

As shown in **Figure 9**, the farnesol-like sesquiterpenoids and natural products nerolidol and bisabolol, which are weakly active in YPD but potent piericidin A suppressors in YPA. Farnesol itself is active in YPA only, as is the natural product parthenolide, which contains a farnesol group in a closed ring conformation instead of the floppy aliphatic conformation of nerolidol and bisabolol (**Figure 9A**). Interestingly, the structurally related natural product 3-hydroxy-4-(succin-2-yl)-caryolane delta-lactone has the opposite profile of parthenolide: it is active in YPD but not in YPA. Analogs of nerolidol and bisabolol that lack a complete farnesol group like linalool and citronellal are inactive, as are closed-ring analogs like menthol and limonene (**Figure 9A**). Piericidin A suppression appears specific to farnesol and farnesol-like sesquiterpenoids because geranylgeraniol is inactive.

**Figure 9.**
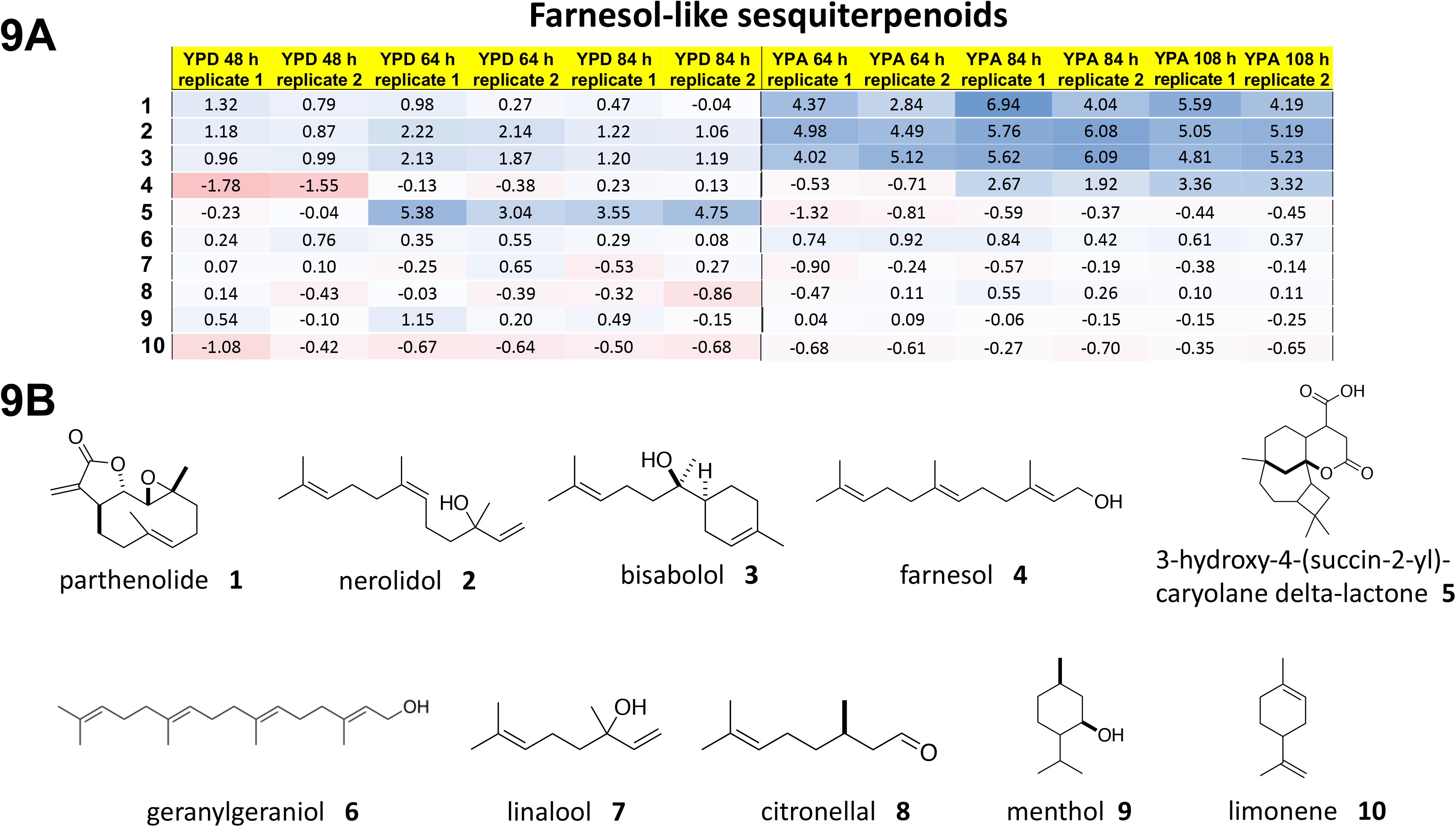
Farnesol-like sesquiterpenoids are piericidin A suppressors. (A) Heat map of Z-scores. The higher the Z-score, the bluer the cell. The lower the Z-score, the redder the cell. A Z-score of zero is a white cell. Columns are screening conditions and timepoints of absorbance measurements. Rows are compounds. (B) Chemical structures showing structure-activity relationships.

There are three smaller classes of piericidin A suppressors that contain one or two active test compounds. For example, the gedunin series (**Figure S3**), proton pump inhibitors (**Figure S4**), and celastrols (**Figure S5**). Gedunins are plant-based natural products and triterpenoids with a furanolactone core scaffold. Among the gedunin series of 10 analogs, most single-site substitutions do not appear to be tolerated. Among the six proton pump inhibitors in the Microsource Spectrum library, only lansoprazole is active. Interestingly, the enantiomer of lansoprazole is not active. Celastrol and dihydrocelastrol are potent and fast-acting piericidin A suppressors in both YPD and YPA.

### Rapamycin counter-screen

Because a piericidin A suppressor screen has never been performed or at least published, it is unclear which of the piericidin A suppressors described herein are selective for piercidin A. Based on the work of my graduate thesis project (Sarkar *et al.*, 2007), I performed a rapamycin chemical modifier screen with the Microsource Spectrum collection as a counter-screen to identify which (if any) piericidin A suppressors are also rapamycin suppressors. The results of this counter-screen are shown in **Figure S6**. Only three test compounds are suppressors of rapamycin: alpha-mangostin, clofoctol and closantel. That number increases to four suppressors of rapamycin when a test compound with discordant replicates is included: rafoxanide. As expected, none of these compounds were active in the piericidin A drug repurposing screen – and none of these compounds have been previously described as suppressors of rapamycin. Reassuringly, closantel, rafoxanide and clofoctol share a core scaffold (**Figure S7**).

Those three compounds, along with two other structural analogs that are in the Microsource Spectrum collection but inactive in the primary screen, were reordered and retested in dose-response experiments. Alpha-mangostin was reordered as was its close analog gamma-mangostin, which differs by a single methyl substitution (**Figure S7**). Dose-response experiments showed conclusively that all four primary screen hits are real. In fact, alpha-mangostin, gamma-mangostin, closantel and rafoxanide all have sub-micromolar potency as suppressors of rapamycin (**Figure S8**). Oxyclozanide and diclaruzil, which are both inactive in the primary screen are completely inactive upon retesting. These results instill confidence in the piericidin A suppressors as selective repurposing candidates and suggest that inactivity in the primary screen is predictive of inactivity in secondary dose-response retests.

## Discussion

In summary, I describe the results of the first-ever drug repurposing screen for mitochondrial diseases, specifically complex I deficiencies including Leigh syndrome, using *Yarrowia lipolytica* yeast cells treated with the complex I inhibitor piericidin A. 24 statistically significant novel suppressors of piericidin A were identified. These suppressors include both FDA approved drugs and generally recognized as safe (GRAS) compounds. A counter-screen of the Microsource Spectrum collection for rapamycin suppressors demonstrates that all 24 piericidin A suppressors are specific. Structure-activity relationships analysis revealed at least four structural classes though it is not clear if these structural classes map to distinct pharmacological classes. Estrogens and estrogen receptor agonists comprise the first and largest group of piericidin A suppressors that are active in both media conditions and at multiple timepoints. Calcium channel blockers, including both dihydropyridines and non-dihydropyridines, comprise the second group of piericidin A suppressors, and are more potent and faster acting in YPA than in YPD. Phenothiazines comprise the third group of piericidin A suppressors. Farnesol-like sesquiterpenoids comprise the fourth group of piericidin A suppressors. There are also three smaller groups of piericidin A suppressors to consider. Overall, the primary screen had a hit rate consistent with previous model organism-based drug repurposing efforts (Rodriguez *et al.*, 2018; Lao *et al.*, 2019; Iyer *et al.*, 2019a; Iyer *et al.*, 2019b).

The estrogens are the most interesting class of piericidin A suppressor because the synthetic estrogen receptor agonist hexestrol (Solmssen, 1945) is an overlapping hit between this *Yarrowia lipoytica* drug repurposing screen and the *Schizosaccharomyces pombe* drug repurposing screen by Delerue *et al.* The authors of that study postulated that hexestrol inhibits mitochondrial fission resulting in suppression of the mitochondrial fragmentation phenotype in a *msp1*^*P300S*^ mutant, and resulting in a mitochondrial hyper-filamentation phenotype in wildtype *Schizosaccharomyces pombe* cells (Delerue *et al.*, 2019). At this point in time, one can only speculate as to the exact mechanism of action of hexestrol. Estrogens have been reported to augment mitochondrial function but their inherent polypharmacology makes pinpointing drug targets difficult (Simpkins & Dykens, 2008). Because *Yarrowia lipolytica* does not encode any obvious orthologs of estrogen receptors, it is possible that hexestrol as well as natural and synthetic estrogens interact with mitochondrial membranes in such a way as to affect membrane biophysical properties, including the mitochondrial membrane potential, which in turn affect mitochondrial protein complexes embedded in the mitochondrial membrane (Torres *et al.*, 2018). The increased probability of onset of Parkinson’s disease in post-menopausal women (Wooten *et al.*, 2004) and the higher incidence of Leber’s hereditary optic neuropathy (LHON) in males versus females (Tonska *et al.*, 2010) could be explained by mitoprotective effects of estrogens. If mitochondrial membranes are targets for estrogens, one also cannot rule out antioxidant effects and glutathione-dependent quenching of reactive oxygen species (ROS), or effects on mitochondrial ion homeostasis, in particular calcium channels.

The dihydropyridine and non-dihydropyridine calcium channel blockers and the farnesol-like sesquiterpenoids may both be affecting mitochondrial calcium homeostasis. Phenothiazines have polypharmacology but could also be affecting ion homeostasis (Barygin *et al.*, 2017). Elucidation of the mechanism of action of the gedunins, celastrols, and proton pump inhibitor lansoprazole as piericidin A suppressors is not possible in the absence of additional experiments to untangle polypharmacology. However, each of those compound classes have been shown to have antioxidant effects and prevent lipid peroxidation (Rai *et al.*, 2011).

Several important caveats to keep in mind and logical next steps. First, the SAR analysis presented herein is based on the primary screening data, which involved just a single concentration of test compound (25μM). Inactivity of a test compound in the primary screen does not necessarily mean that said compound is inactive at concentrations higher than 25μM, though cytotoxicity is increasingly a concern above 25μM. Second, none of the piericidin A suppressors have been validated in a genetic model of mitochondrial disease whether in *Yarrowia lipolytica* or any other model system. NDUF8A is a nuclear gene that encodes an assembly factor specific to complex I (Stroud *et al.*, 2016). *Yarrowia lipolytica* has an ortholog of *NDUFA8*; truncation mutants are, as expected, inviable (Dr Mark Blenner, personal communication). As has been done for another nuclear encoded assembly factor gene in *Yarrowia lipolytica* (Gerber *et al.*, 2017), hypomorphic *NDUFA8* missense mutants are being generated now in collaboration with a lab at Clemson University so that the piericidin A suppressors described herein will be tested for their ability to rescue the growth defects of this mutant. Once validated in yeast and patient derived cells (fibroblasts, followed by iPSCs), piericidin A suppressors can be further validated in complex I deficient worms (Polyak *et al.*, 2018), flies (Cabirol-Pol *et al.*, 2018) and zebrafish (Pinho *et al.*, 2013). Cross-species validated piericidin A suppressors that are FDA approved drugs have a clear path to the clinic, starting in so called N-of-1 trials.

## Acknowledgements and Funding

Without the support of the following individuals and labs this project would not have been possible. First and foremost, I acknowledge Radical Investments, LLC, the venture capital firm of entrepreneur and investor Mark Cuban, as the primary funding source. I acknowledge Collaborations Pharma, Inc based in Raleigh, North Carolina for allowing me to conduct these experiments in their lab space during the Spring of 2019, specifically Dr. Sean Ekins, Thomas Lane, and Dr. Ana Puhl Rubio. I acknowledge Brianne Vignero from Dr. Daniel Burke’s lab at North Carolina State University for preparing sterile media and overnight cultures. I acknowledge Dr Patrick Gibney at Cornell University for performing piericidin A dose-response experiments to determine the IC50 doses in glucose-containing media and in acetate-containing media. I acknowledge Vidya Seshadri and Dr. So Young Kim at Duke University Medical School for use of their Labcyte Echo550. I acknowledge Dr. Maitreya Dunham at the University of Washington for generously providing the BY prototroph strain. I acknowledge an ongoing collaboration with Dr Mark Blenner at Clemson University. Finally, I acknowledge Tom Ruginis from HappiLabs, Inc for ordering supplies and reagents for this project.

---

**Figure S1**. Non-estrogenic steroids are not piericidin A suppressors. (A) Heat map of Z-scores. The higher the Z-score, the bluer the cell. The lower the Z-score, the redder the cell. A Z-score of zero is a white cell. Columns are screening conditions and timepoints of absorbance measurements. Rows are compounds. (B) Chemical structures showing structure-activity relationships.

**Figure S2**. Non-dihydropyridine calcium channel blockers are piericidin A suppressors. (A) Heat map of Z-scores. The higher the Z-score, the bluer the cell. The lower the Z-score, the redder the cell. A Z-score of zero is a white cell. Columns are screening conditions and timepoints of absorbance measurements. Rows are compounds. (B) Chemical structures showing structure-activity relationships.

**Figure S3**. Gedunins are piericidin A suppressors. (A) Heat map of Z-scores. The higher the Z-score, the bluer the cell. The lower the Z-score, the redder the cell. A Z-score of zero is a white cell. Columns are screening conditions and timepoints of absorbance measurements. Rows are compounds. (B) Chemical structures showing structure-activity relationships.

**Figure S4**. The proton pump inhibitor lansoprazole is a piericidin A suppressor. (A) Heat map of Z-scores. The higher the Z-score, the bluer the cell. The lower the Z-score, the redder the cell. A Z-score of zero is a white cell. Columns are screening conditions and timepoints of absorbance measurements. Rows are compounds. (B) Chemical structures showing structure-activity relationships.

**Figure S5**. Celastrol and dihydrocelastrol are piericidin A suppressors. (A) Heat map of Z-scores. The higher the Z-score, the bluer the cell. The lower the Z-score, the redder the cell. A Z-score of zero is a white cell. Columns are screening conditions and timepoints of absorbance measurements. Rows are compounds. (B) Chemical structures showing structure-activity relationships.

**Figure S6**. Rapamycin counter-screen of the Microsource Spectrum library. Yeast cells were treated with 50nM rapamycin and test compounds in 384-well plates and incubated for four days at room temperature without shaking. Gray dots are test compounds. Replicate one and replicate two are plotted against each other. X-axis and y-axis denotes OD_600_ absorbance measurements.

**Figure S7**. Structure-activity relationships of novel rapamycin suppressors. The chemical structures of the rapamycin suppressors clofoctol, closantel and rafoxanide alongside inactive analogs oxyclozanide and diclazuril; and the rapamycin suppressor alpha-mangostin alongside its active analog gamma-mangostin. Shaded regions indicate sites of substitution.

**Figure S8**. Dose-response experiments show that activity (or inactivity) in the primary drug screen is predictive of activity (or inactivity) in multi-concentration-point retests. Two-fold titrations spanning three orders of magnitude from 100μM down to 120nM are denoted on the x-axis. OD600 absorbance measurements are denoted on the y-axis.

