## Supplemental Figures for "Drug repurposing for mitochondrial diseases using a pharmacological model of complex I deficiency in the yeast *Yarrowia lipolytica*"

# S1

#### Steroids

| | YPD 48 h replicate 1 | YPD 48 h replicate 2 | YPD 64 h replicate 1 | YPD 64 h replicate 2 | YPD 84 h replicate 1 | YPD 84 h replicate 2 | YPA 64 h replicate 1 | YPA 64 h replicate 2 | YPA 84 h replicate 1 | YPA 84 h replicate 2 | YPA 108 h replicate 1 | YPA 108 h replicate 2 |
|---|---|---|---|---|---|---|---|---|---|---|---|---|
| 1 | -1.00 | -0.09 | 0.66 | 0.32 | 3.46 | 0.79 | -0.27 | -0.52 | 0.18 | 0.06 | 0.21 | 0.23 |
| 2 | -0.24 | -0.60 | -0.30 | -0.10 | 0.79 | 2.64 | -0.12 | -0.15 | 0.42 | 0.38 | 0.23 | 1.29 |
| 3 | 4.52 | 4.68 | 4.23 | 4.28 | 2.57 | 2.65 | 1.61 | 1.03 | 1.95 | 1.71 | 2.04 | 2.11 |
| 4 | 2.42 | 4.85 | 4.71 | 3.30 | 3.11 | 2.13 | -0.61 | 0.16 | 0.29 | 1.60 | 0.66 | 2.08 |
| 5 | 0.31 | -0.45 | 1.85 | 1.53 | 2.40 | 1.23 | -1.41 | -0.96 | -0.84 | 0.11 | -0.80 | 0.77 |
| 6 | 0.80 | 0.21 | 0.24 | 0.40 | 0.33 | 0.44 | 0.31 | -0.14 | 0.15 | -0.26 | -0.04 | -0.44 |
| 7 | 0.22 | -0.56 | -0.25 | -1.25 | -0.20 | -0.77 | 0.82 | -0.48 | -0.20 | -0.46 | -0.06 | -0.34 |
| 8 | -0.07 | 0.02 | -0.30 | -0.36 | -0.25 | -0.43 | -0.34 | -0.64 | -0.51 | -0.83 | -0.68 | -0.83 |
| 9 | 1.37 | 2.20 | 2.05 | 2.85 | 1.23 | 1.63 | 1.21 | 2.54 | 3.06 | 3.80 | 3.07 | 3.86 |
| 10 | 1.62 | 0.98 | 1.90 | 1.47 | 0.97 | 0.64 | 0.88 | 0.80 | 0.53 | 0.91 | 1.39 | 1.48 |

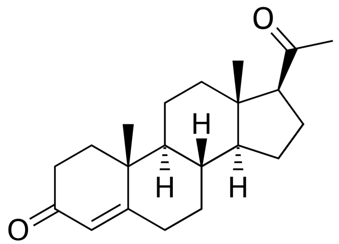

progesterone **1**

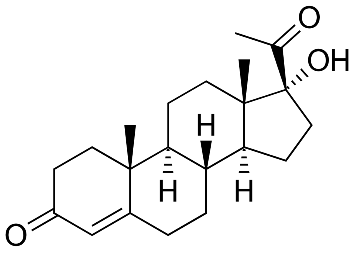

hydroxyprogesterone **2**

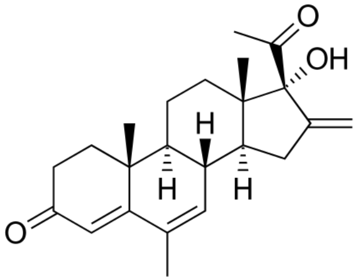

melengestrol **3**

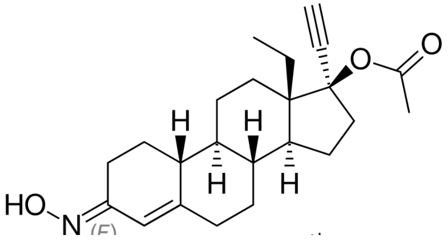

norgestimate **4**

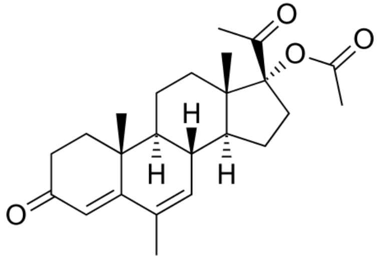

megestrol acetate **5**

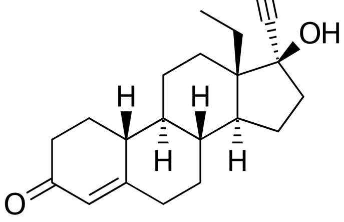

levonorgesterel **6**

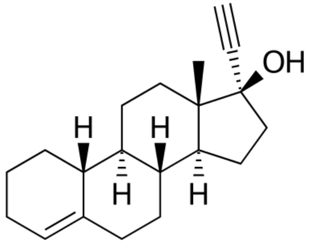

lynestrenol **7**

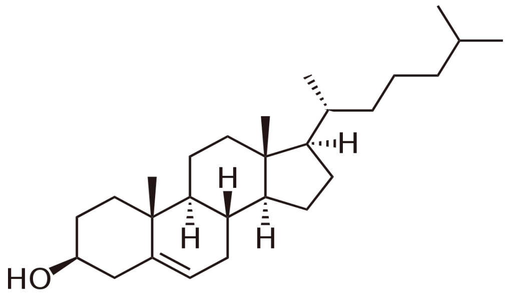

cholesterol **8**

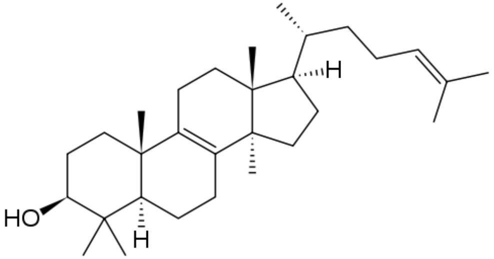

lanosterol **9**

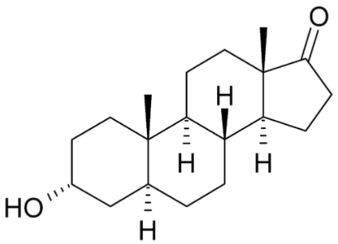

androsterone **10**

# S2

#### Calcium channel blockers

| | YPD 48 h replicate 1 | YPD 48 h replicate 2 | YPD 64 h replicate 1 | YPD 64 h replicate 2 | YPD 84 h replicate 1 | YPD 84 h replicate 2 | YPA 64 h replicate 1 | YPA 64 h replicate 2 | YPA 84 h replicate 1 | YPA 84 h replicate 2 | YPA 108 h replicate 1 | YPA 108 h replicate 2 |
|---|---|---|---|---|---|---|---|---|---|---|---|---|
| **1** | -2.19 | -2.67 | -2.34 | -2.74 | -2.44 | -2.49 | 3.62 | 3.86 | 4.13 | 4.50 | 3.29 | 3.67 |
| **2** | -0.16 | -0.50 | -0.64 | -0.95 | -1.06 | -1.29 | 1.10 | 2.15 | 2.63 | 5.15 | 3.73 | 5.50 |
| **3** | 0.80 | 0.87 | 1.42 | 1.54 | 0.65 | 0.96 | 2.34 | 3.33 | 4.26 | 5.29 | 3.92 | 4.70 |
| **4** | 0.43 | 0.00 | 0.43 | 0.64 | 0.50 | 0.54 | 0.67 | 0.09 | 0.11 | 0.14 | 0.14 | -0.03 |
| **5** | 1.83 | 0.98 | 0.32 | -0.27 | 0.05 | -0.18 | 2.24 | 1.74 | 1.39 | 1.36 | 0.99 | 1.02 |

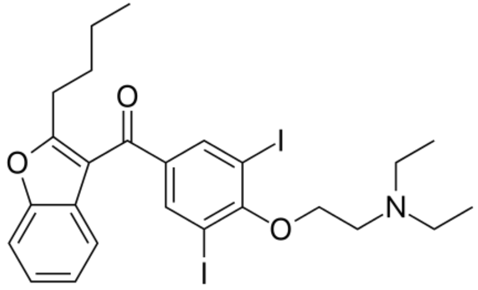


amiodarone **1**

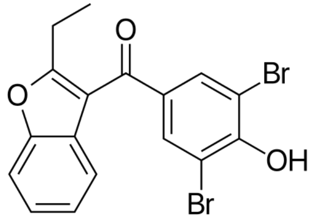


benzbromarone **2**

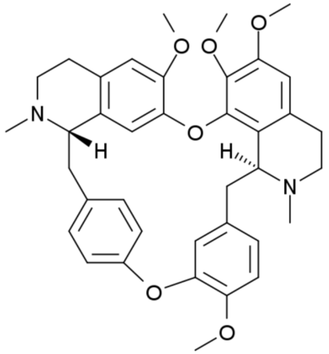


tetrandrine **3**

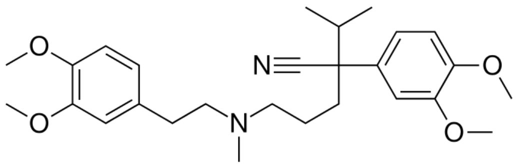


verapamil **4**

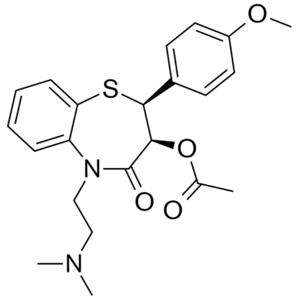


diltiazem **5**

# S3

### Gedunins

| | YPD 48 h replicate 1 | YPD 48 h replicate 2 | YPD 64 h replicate 1 | YPD 64 h replicate 2 | YPD 84 h replicate 1 | YPD 84 h replicate 2 | YPA 64 h replicate 1 | YPA 64 h replicate 2 | YPA 84 h replicate 1 | YPA 84 h replicate 2 | YPA 108 h replicate 1 | YPA 108 h replicate 2 |
|---|---|---|---|---|---|---|---|---|---|---|---|---|
| **1** | 4.16 | 4.37 | 4.19 | 4.04 | 2.14 | 2.09 | 6.22 | 6.27 | 6.81 | 6.95 | 5.98 | 6.32 |
| **2** | 2.38 | 1.50 | 3.88 | 2.10 | 2.11 | 0.99 | 2.56 | 1.86 | 6.73 | 4.83 | 6.03 | 5.29 |
| **3** | 5.08 | 2.38 | 4.78 | 2.91 | 2.33 | 1.72 | 2.32 | 2.22 | 2.68 | 3.24 | 3.33 | 3.54 |
| **4** | 1.29 | 0.93 | 1.96 | 1.38 | 1.05 | 0.68 | 0.84 | 1.26 | 2.14 | 1.16 | 2.59 | 2.11 |
| **5** | 1.11 | 3.07 | 3.34 | 3.46 | 1.84 | 2.34 | 0.07 | 0.27 | 1.10 | 1.36 | 4.08 | 2.24 |
| **6** | 0.14 | -0.03 | 1.24 | 1.01 | 0.69 | 0.45 | -0.53 | -0.30 | 0.49 | 0.14 | 1.49 | 0.94 |
| **7** | -0.40 | 0.09 | 1.25 | 0.60 | 0.88 | 0.37 | -0.74 | -1.37 | -0.47 | -1.37 | 0.64 | -0.66 |
| **8** | -0.31 | -0.41 | 0.03 | -0.09 | -0.06 | -0.34 | -0.52 | -0.31 | 0.03 | -0.11 | 0.11 | -0.06 |
| **9** | -0.73 | -1.07 | -0.49 | -0.89 | -0.50 | -0.65 | -0.76 | -0.91 | -0.44 | -0.67 | -0.65 | -0.77 |
| **10** | 0.50 | 0.84 | 0.17 | 1.09 | -0.13 | 0.71 | 0.61 | 1.00 | 0.57 | 0.59 | 0.47 | 0.41 |

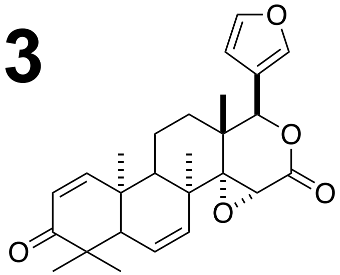

7-desacetoxy-6,7-dehydrogedunin **1**

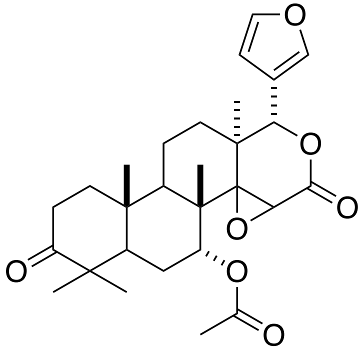

dihydrogedunin **2**

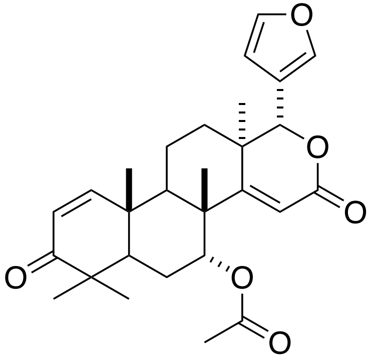

deoxgedunin **4**

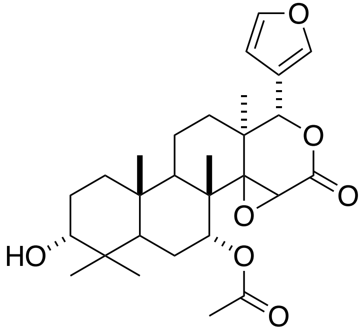

gedunol **5**

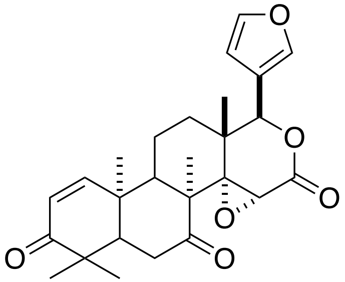

3alpha-acetoxydihydro deoxygedunin **6**

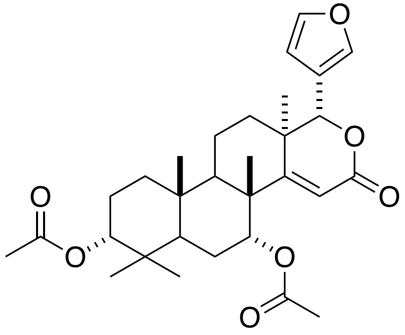

deacetoxy-7-oxogedunin **7**

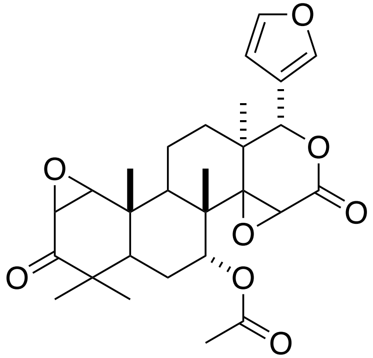

epoxygedunin **3**

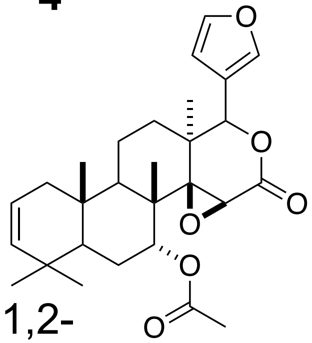

3,4-dehydro-1,2-dihydro-3-desoxo-gedunin **8**

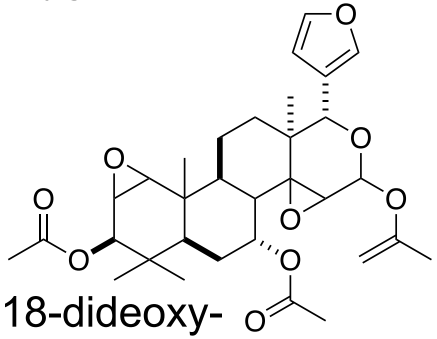

1,2-epoxy-3,18-dideoxy-3beta,18-diacetyoxygedunin **9**

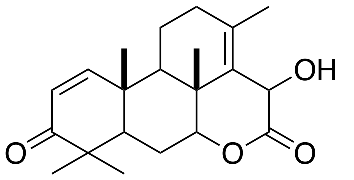

merogedunin **10**

**S4**

### Proton pump inhibitors

| | YPD 48 h replicate 1 | YPD 48 h replicate 2 | YPD 64 h replicate 1 | YPD 64 h replicate 2 | YPD 84 h replicate 1 | YPD 84 h replicate 2 | YPA 64 h replicate 1 | YPA 64 h replicate 2 | YPA 84 h replicate 1 | YPA 84 h replicate 2 | YPA 108 h replicate 1 | YPA 108 h replicate 2 |
|---|---|---|---|---|---|---|---|---|---|---|---|---|
| **1** | 1.01 | 0.27 | 3.91 | 1.65 | 2.38 | 5.48 | -0.79 | -0.07 | 0.49 | 0.15 | 3.91 | 2.12 |
| **2** | 0.47 | 0.04 | 0.30 | 0.48 | 0.57 | 0.30 | 0.28 | 0.79 | 0.43 | 0.24 | 0.57 | 0.46 |
| **3** | 0.33 | -0.29 | 0.35 | 0.17 | 0.41 | 0.35 | 0.50 | -0.15 | 0.22 | -0.22 | 0.69 | 0.31 |
| **4** | 1.50 | 0.84 | 1.35 | 0.35 | 0.99 | 0.56 | 1.38 | 1.13 | 0.67 | 0.90 | 0.66 | 0.77 |
| **5** | -0.73 | -0.39 | -1.10 | 0.51 | -0.80 | 0.58 | 0.10 | -0.21 | 0.26 | 0.09 | 0.22 | 0.64 |
| **6** | -0.21 | -0.54 | -0.54 | -0.04 | -0.30 | -0.23 | -0.50 | -0.66 | -0.73 | -0.48 | -0.53 | -0.48 |

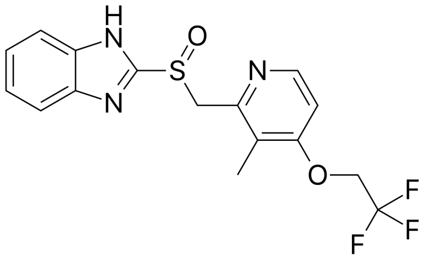
lansoprazole **1**

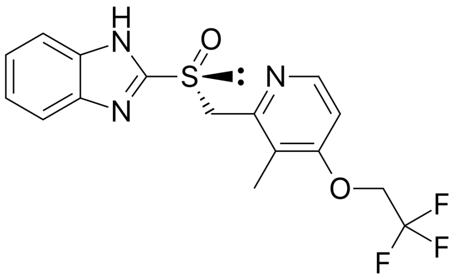
dezlansoprazole **2**

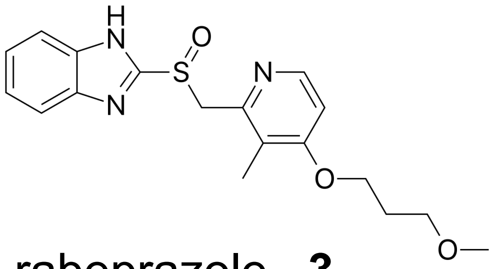
rabeprazole **3**

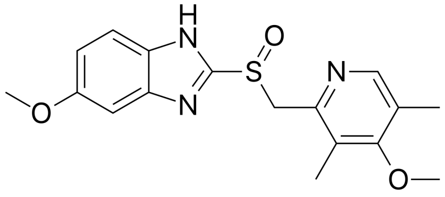
omeprazole **4**

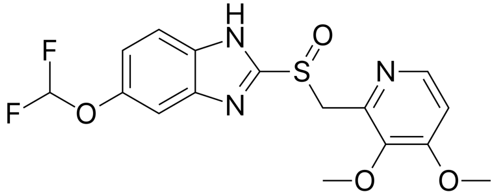
pantoprazole **5**

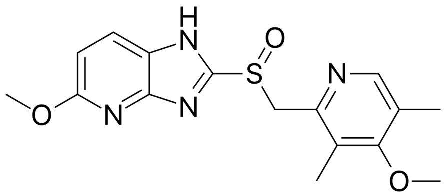
tenatoprazole **6**

# S5

#### Celastrols

| | YPD 48 h replicate 1 | YPD 48 h replicate 2 | YPD 64 h replicate 1 | YPD 64 h replicate 2 | YPD 84 h replicate 1 | YPD 84 h replicate 2 | YPA 64 h replicate 1 | YPA 64 h replicate 2 | YPA 84 h replicate 1 | YPA 84 h replicate 2 | YPA 108 h replicate 1 | YPA 108 h replicate 2 |
|---|---|---|---|---|---|---|---|---|---|---|---|---|
| **1** | 2.33 | 0.84 | 2.55 | 1.54 | 6.24 | 0.79 | 2.90 | 3.26 | 5.82 | 4.89 | 5.20 | 4.98 |
| **2** | 4.80 | 4.48 | 3.80 | 3.46 | 1.87 | 1.65 | 2.55 | 2.27 | 3.64 | 2.98 | 3.97 | 3.63 |

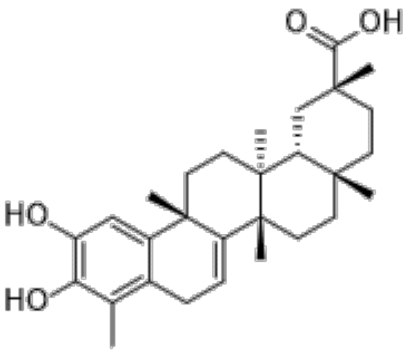


dihydrocelastrol **1**

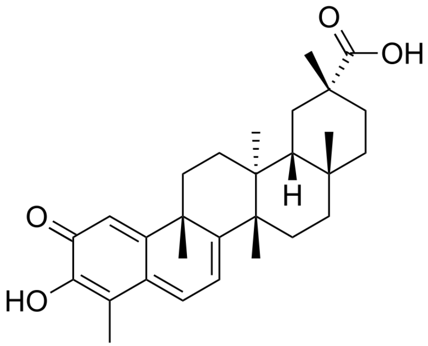


celastrol **2**

# S6

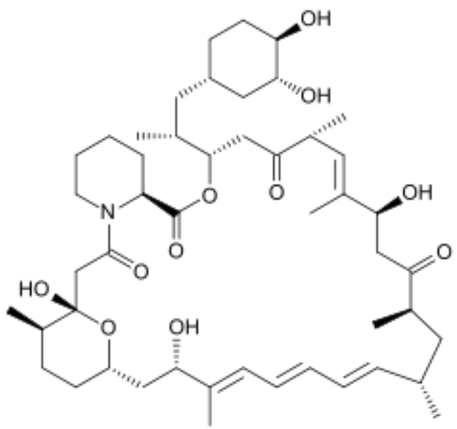


50nM rapamycin
YPD 96 h

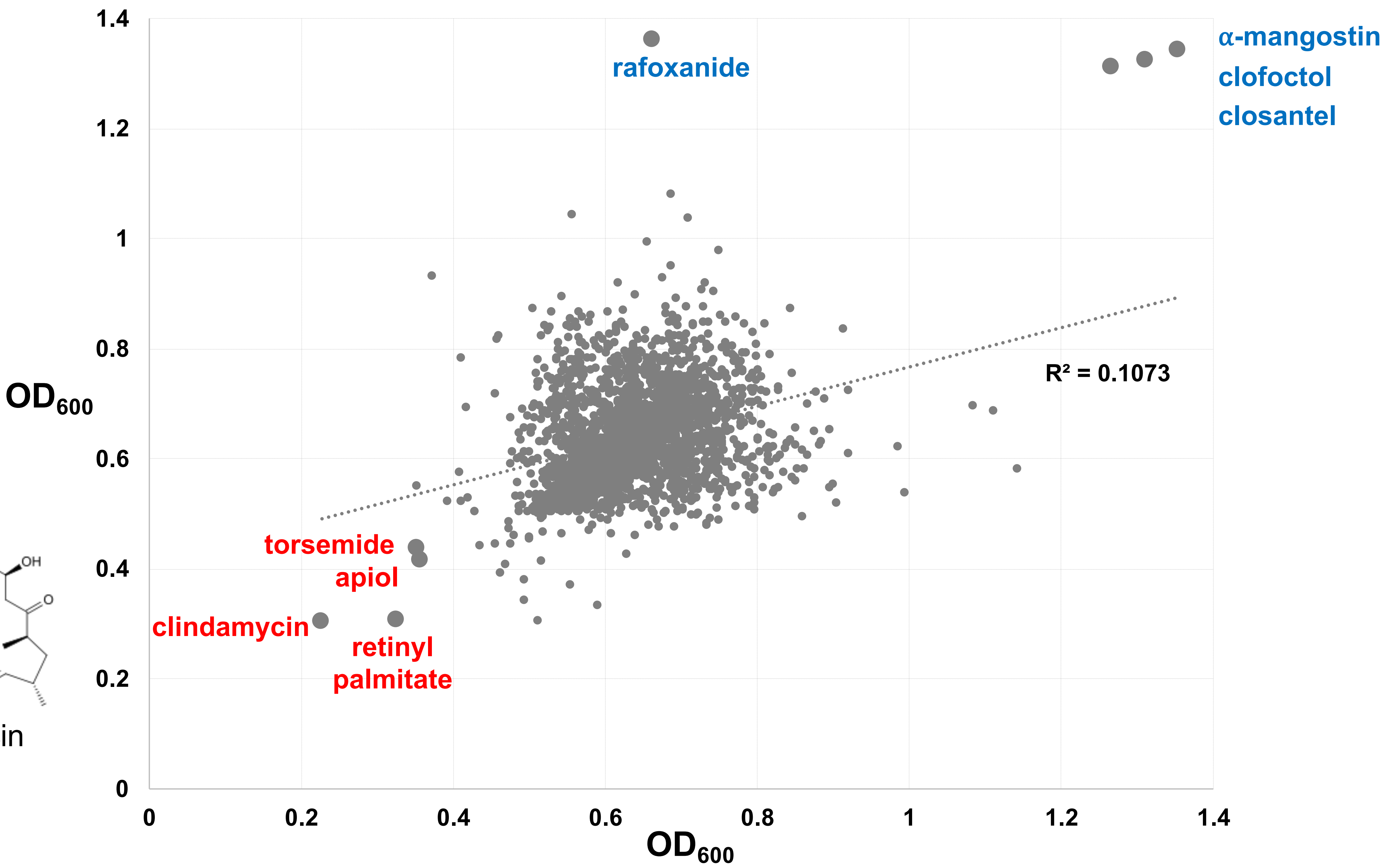

# S7

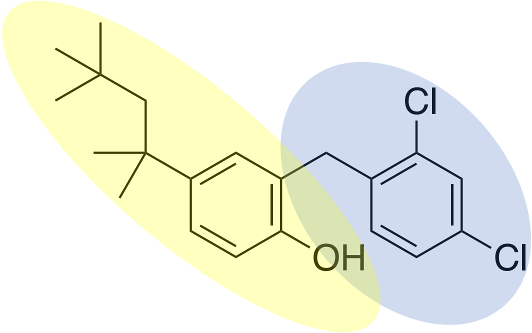

clofoctol

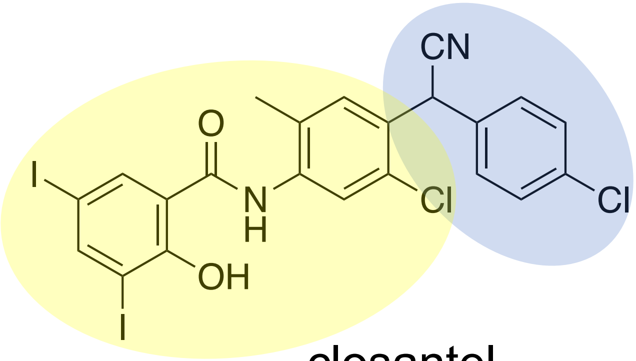

closantel

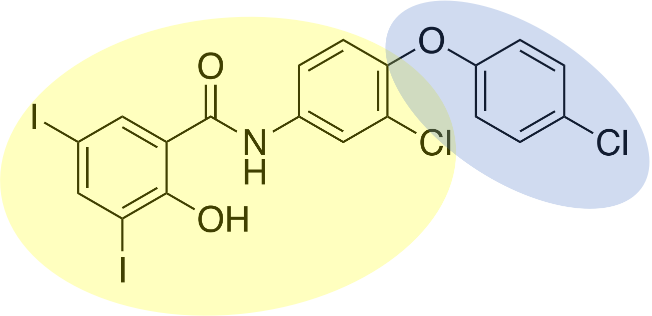

rafoxanide

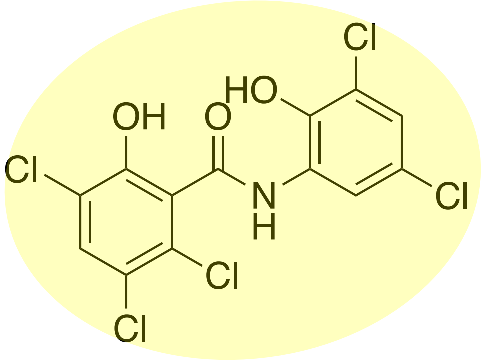

oxyclozanide

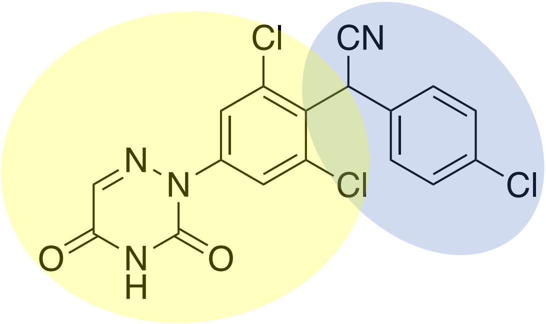

diclazuril

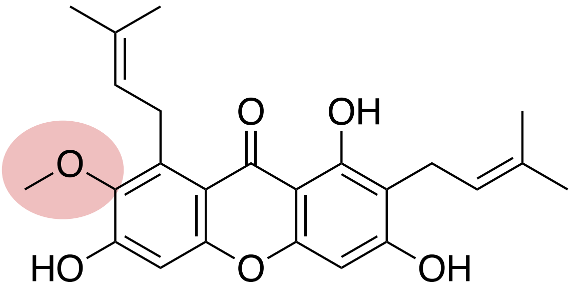

α-mangostin

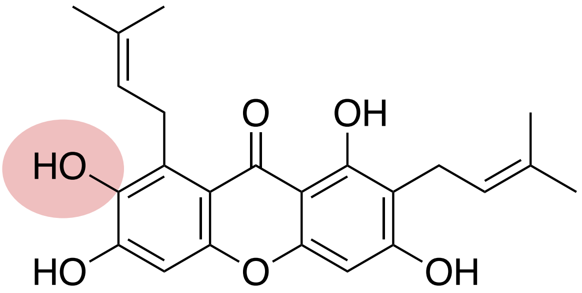

γ-mangostin

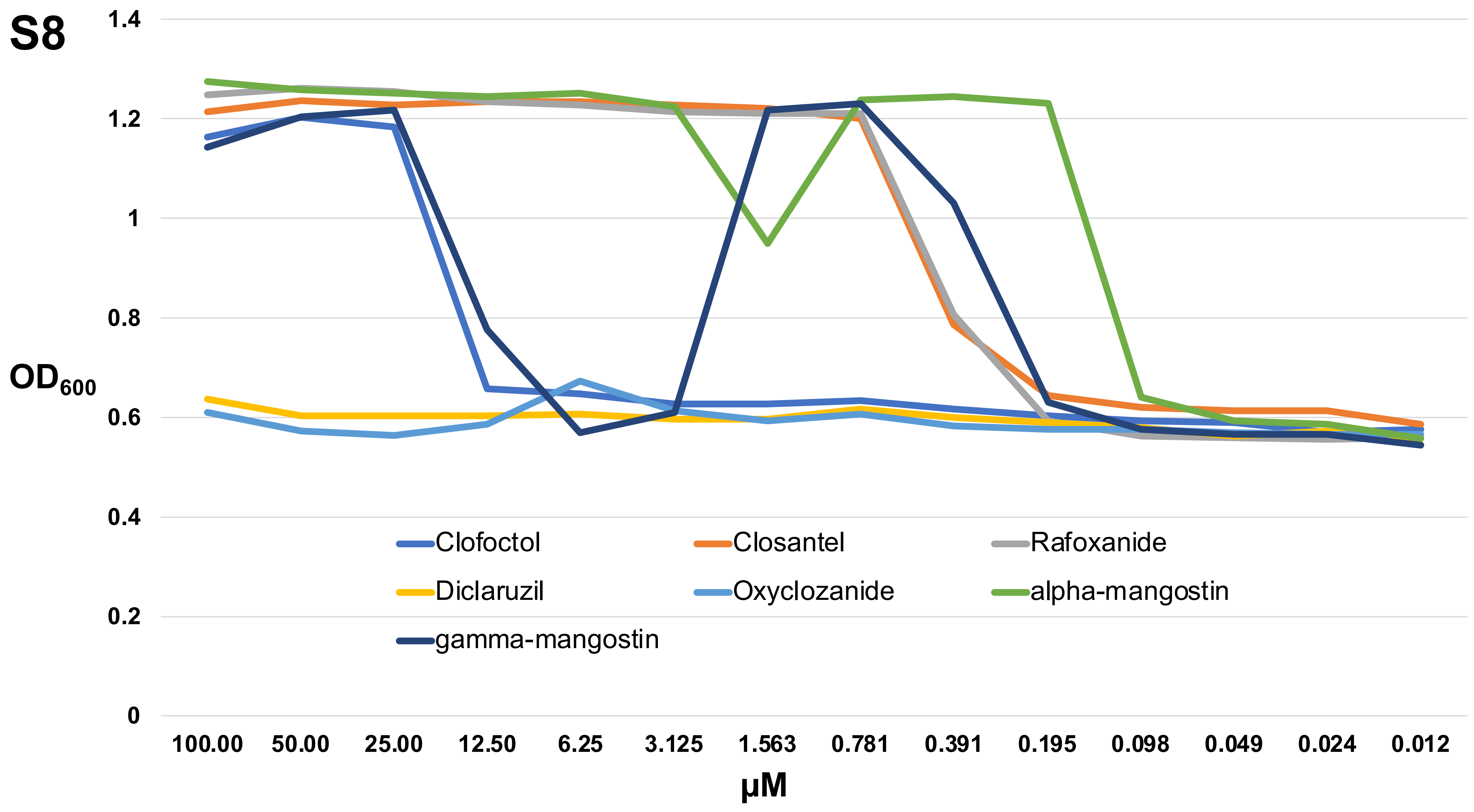
S8
1.4
1.2
1
0.8
0.6
0.4
0.2
0
OD600
Clofoctol
Closantel
Rafoxanide
Diclaruzil
Oxyclozanide
alpha-mangostin
gamma-mangostin
100.00
50.00
25.00
12.50
6.25
3.125
1.563
0.781
0.391
0.195
0.098
0.049
0.024
0.012
µM
